# Dynamic control of proinflammatory cytokines Il-1β and Tnf-α by macrophages is necessary for functional spinal cord regeneration in zebrafish

**DOI:** 10.1101/332197

**Authors:** Themistoklis M. Tsarouchas, Daniel Wehner, Leonardo Cavone, Tahimina Munir, Marcus Keatinge, Marvin Lambertus, Anna Underhill, Thomas Barrett, Elias Kassapis, Nikolay Ogryzko, Yi Feng, Tjakko J. van Ham, Thomas Becker, Catherina G. Becker

**Affiliations:** Centre for Discovery Brain Sciences, University of Edinburgh, The Chancellor’s Building, 49 Little France Crescent, Edinburgh, EH16 4SB, UK; MRC Centre for Inflammation Research, Queen’s Medical Research Institute, University of Edinburgh, Edinburgh, EH16 4TJ, UK; Department of Clinical Genetics, Erasmus University Medical Center, Wytemaweg 80, 3015 CN Rotterdam, the Netherlands

## Abstract

Spinal cord injury leads to a massive response of innate immune cells (microglia, macrophages, neutrophils) both, in non-regenerating mammals and in successfully regenerating zebrafish, but the role of these immune cells in functional spinal cord regeneration in zebrafish has not been addressed. Here we show that inhibiting inflammation reduces and promoting it accelerates axonal regeneration in larval zebrafish. Mutant analyses show that peripheral macrophages, but not neutrophils or microglia, are necessary and sufficient for full regeneration. Macrophage-less *irf8* mutants show prolonged inflammation with elevated levels of Il-1β and Tnf-α. Decreasing Il-1β levels or number of Il-1β^+^ neutrophils rescues functional regeneration in *irf8* mutants. However, during early regeneration, interference with Il-1β function impairs regeneration in *irf8* and wildtype animals. Inhibiting Tnf-α does not rescue axonal growth in *irf8* mutants, but impairs it in wildtype animals, indicating a pro-regenerative role of Tnf-α. Hence, inflammation is tightly and dynamically controlled by macrophages to promote functional spinal cord regeneration in zebrafish.

## INTRODUCTION

Zebrafish, in contrast to mammals, are capable of functional spinal cord regeneration after injury. Recovery of swimming function critically depends on regeneration of axonal connections across the complex non-neural injury site ^1,2^. It is therefore important to determine the factors that allow axons to cross the lesion site in zebrafish.

The immune reaction to injury may be a determining factor for regenerative success ^3^. In mammals, a prolonged immune response, consisting of pro-inflammatory macrophages ^4^, microglia cells ^5^ and neutrophils ^6^ together with cytokines release from other cell types, such as endothelial cells, oligodendrocytes or fibroblasts ^7^ contribute to an inflammatory environment that is hostile to axon regeneration. However, activated macrophages can also promote axon regeneration ^8,9^, suggesting complex roles of the immune response after spinal injury.

In zebrafish, we can dissect the roles of all of these cell types in a system that shows a high capacity for axon regeneration and functional spinal cord repair ^10^. Zebrafish possess an innate immune system from early larval stages and develop an adaptive immune system at juvenile stages, both of which are similar to those in mammals ^11^. Indeed, microglia activation has been described in reaction to spinal cord injury in adult ^12,13^ and larval zebrafish ^14^, suggesting important functions of the immune reaction for functional spinal cord regeneration in zebrafish.

Larval zebrafish regenerate more rapidly than adults. Axonal and functional regeneration is observed within 48 hours after an injury in 3 day-old larvae ^1,2^. At the same time, the larval system presents complex tissue interactions that allow us to better understand how axons can cross a non-neural lesion environment. For example, axons encounter Pdgfrb^+^ fibroblast-like cells that deposit regeneration-promoting Col XII in the lesion site in a Wnt-signalling dependent manner ^1^. These cellular and molecular components are present also in the injury sites of adult zebrafish and mammals ^1^. How cells of the immune system contribute to this growth-conducive lesion site environment in zebrafish is unclear.

Here we show that peripheral macrophages control successful axon regeneration by producing pro-regenerative tumour necrosis factor alpha (Tnf-α) and by reducing levels of the pro-inflammatory cytokine interleukin-1 β (Il-1β). While early expression of *il-1β* promotes axon regeneration, prolonged high levels of Il-1β in the macrophage-less *irf8* mutant are detrimental. Preventing formation of Il-1β producing neutrophils or inhibiting excess *il-1β* directly, largely restored the impaired axonal regrowth and recovery of swimming capacity in *irf8* mutants. This indicates that regulation of a single immune system-derived factor, Il-1β, is a major determinant of successful spinal cord regeneration in a complex lesion site environment.

## RESULTS

### The innate immune response coincides with axonal regeneration

We analysed invasion of the injury site by immune cells and other cell types in relation to axonal regrowth in larval zebrafish that underwent complete spinal cord transection at 3 days post-fertilisation (dpf). Axons were present in the injury site by 1day post-lesion (dpl). Axon density in the lesion site was strongly increased by 2 dpl and thereafter plateaued for up to at least 4 dpl (Fig. S1A). This is consistent with previous results showing continuous axon labelling over the lesion site (termed axon bridging) in 80% of animals and recovery of swimming behaviour already by 2 dpl ^14^.

We observed a rapid and massive influx of immune cells, with neutrophils (Mpx^+^) peaking at 2 hours post-lesion (hpl) and macrophages (*mpeg1*:GFP^+^; 4C4^-^) and microglia (mpeg1:GFP^+^; 4C4^+^) accumulating in the lesion site a few hours later and peaking at 2 dpl (Fig. 1A, Movie S1). Myelinating cells (*cldnK*:GFP^*+*^) and endothelial cells (*fli1*:GFP^*+*^) were not abundant in the lesion site during axon regrowth (Fig. S1C,D), in contrast to functionally important *pdgfrb:*GFP^+^ fibroblasts ^1^ that were robustly present in the lesion site at 1 dpl, peaking at 2 dpl (Fig. S1B). This suggests that myelinating cells and endothelial cells are not important for axon bridging. In contrast, the spatio-temporal pattern of immune cell invasion of the injury site suggests a role for the immune system in orchestrating regeneration of axons over the lesion site.

**Fig. 1:**
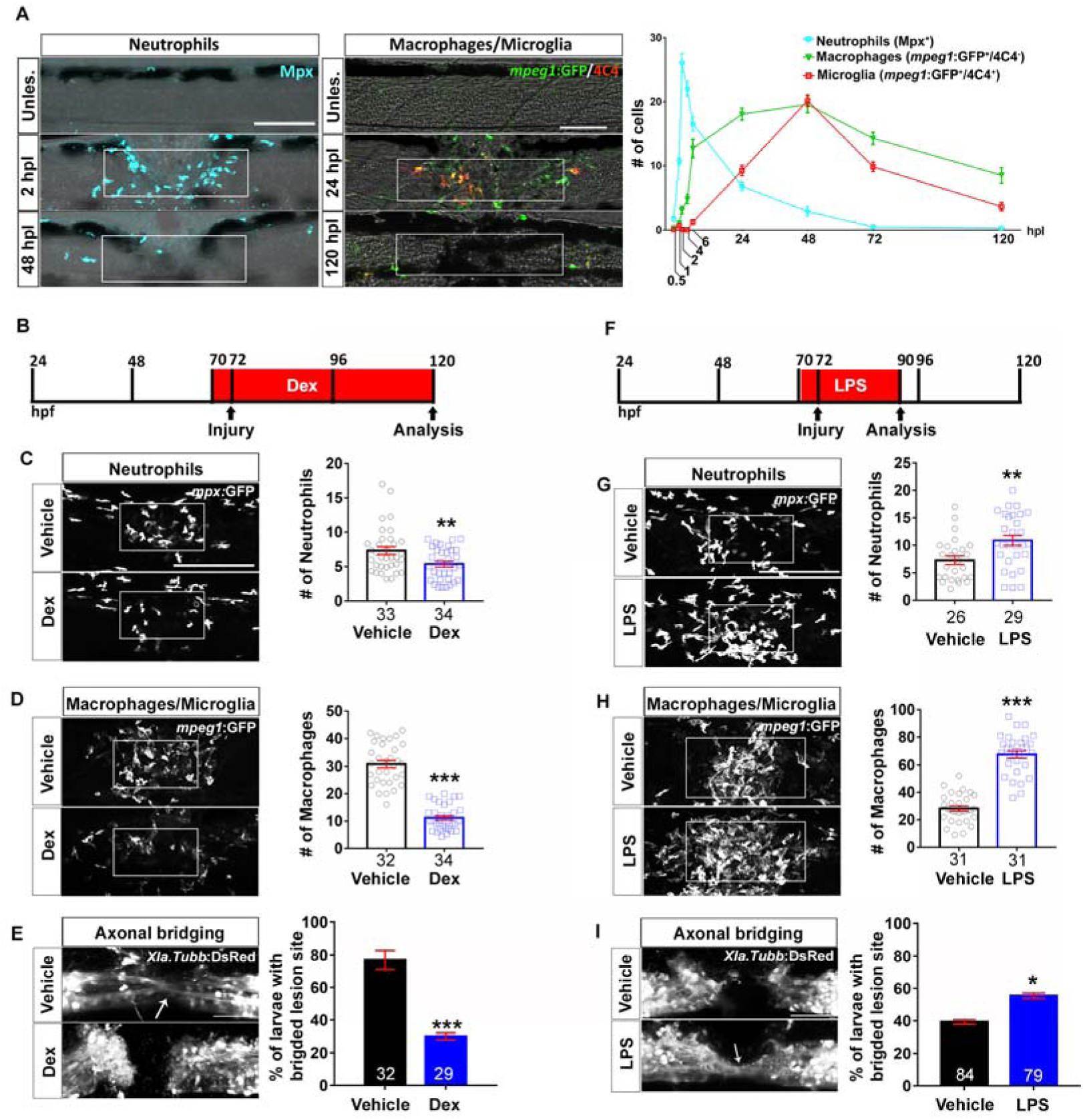
Spinal injury leads to an inflammatory response that promotes axonal regeneration. **A:** Neutrophils, macrophages and microglial cells show different dynamics after injury. Neutrophils (Mpx^+^) accumulate in the injury site very early, peaking at 2hpl. Macrophages (*mpeg1*:GFP^+^/4C4^-^) and microglial cell (*mpeg1*:GFP^+^/4C4^+^) numbers peak at 48hpl. Fluorescence images were projected onto transmitted light images. **B-D:** Incubation with dexamethasone (timeline in B) reduces neutrophil and macrophage numbers (C; *Mann-Whitney U test*: **P<0.01, ***, P<0.001), as well as the proportion of animals with axonal bridging (E; *Fisher’s exact test*: ***P<0.001). **F-H:** Incubation of animals with LPS during early regeneration (timeline in F) increased numbers of neutrophils and macrophages (G, H; *t-test*: **P<0.01, ***, P<0.001), as well as the proportion of animals with axonal bridging (I; *Fisher’s exact test*: *P<0.05). Lateral views of the injury site are shown; rostral is left. Dashed rectangle indicates region of quantification; arrows indicate axonal bridging. *Scale bars*: 50μm; Error bars indicate SEM.

### Immune system activation promotes axonal regeneration

To determine the importance of the immune reaction to a lesion, we inhibited it using the anti-inflammatory synthetic corticosteroid dexamethasone ^14^. This reduced reduced the proportion of larvae exhibiting axon bridging (control: 78%, Dex: 30%; Fig. 1E). Similarly, reducing the immune response, using a well-established combination of morpholinos against *pu.1* and *gcsfr* ^15^, reduced the proportion of larvae with axonal bridges from 81% to 57% (Fig. S2A,B).

For a gain-of-function approach, we enhanced the immune reaction by incubation with bacterial lipopolysaccharides (LPS). This increased the number of neutrophils and macrophages in the lesion site (Fig. 1F-H). To detect a potential accelerating effect on axon regrowth we analysed larvae at 18 hpl, when axon regeneration was incomplete in untreated lesioned animals. This showed an increase in the proportion of larvae with axonal bridges from 41% in wildtype to 60% in LPS-treated animals (Fig. 1I). Hence, immune system activity is necessary for and sufficient to promote axon regeneration across a spinal lesion site.

### Macrophages determine regenerative success

To analyse the role of different immune cell types in the regeneration process, we used mutant analysis. In mutants for the macrophage-lineage determining transcription factor *irf8*, macrophages and microglial cells, but not neutrophils are missing during early stages of development ^16^. These homozygous mutants are adult viable and show no overt developmental aberrations, except for an increased number of neutrophils ^16^. In situ hybridisation for the macrophage and microglia marker *mpeg1* confirmed ample signal in the ventral trunk region of unlesioned larvae and in a spinal lesion site at 2 dpl in wildtype larvae, but complete absence of in situ hybridisation signal in unlesioned and lesioned *irf8* larvae (Fig. 2A).

**Fig. 2:**
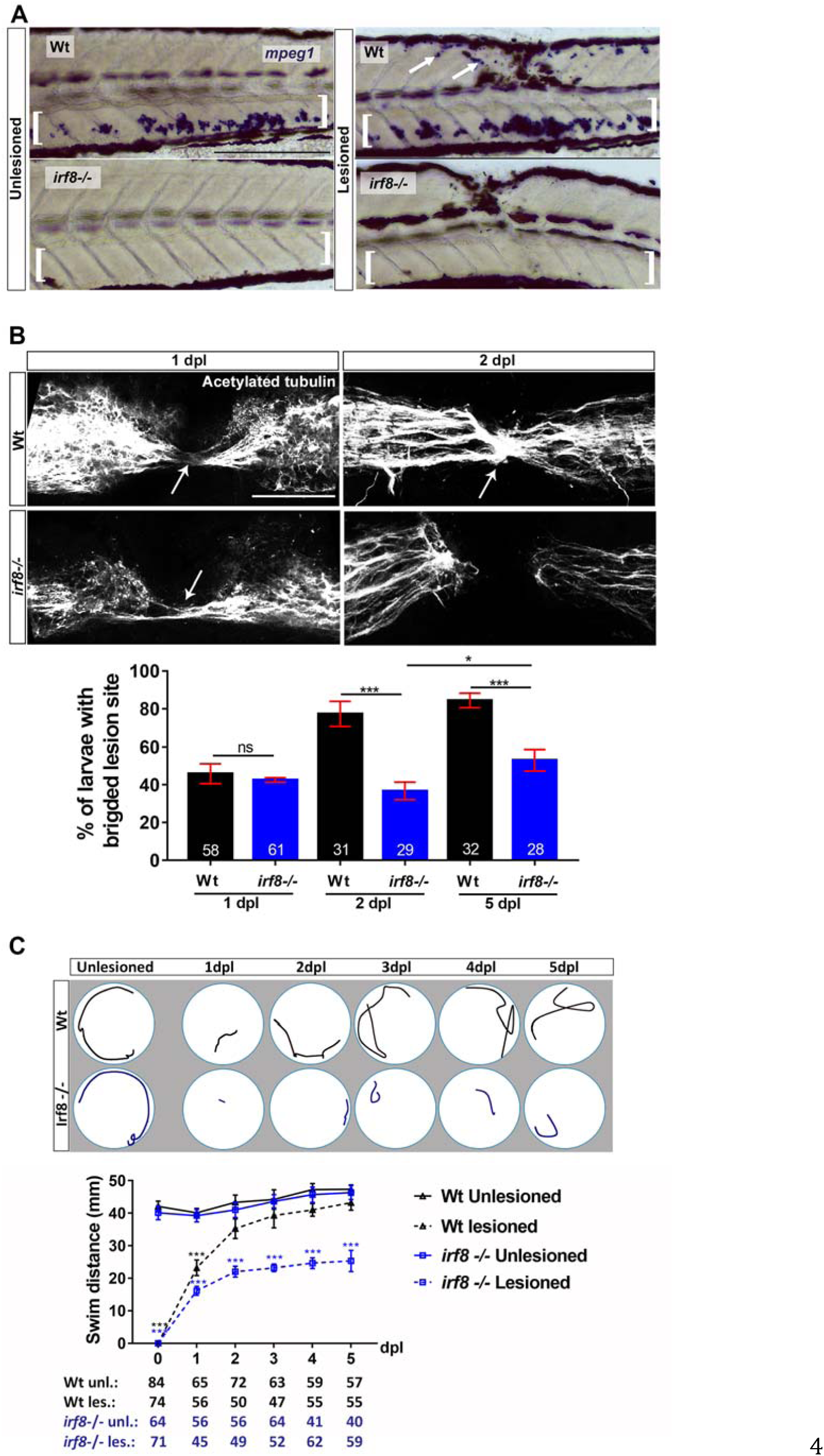
In the *irf8* mutant, axonal regeneration and functional recovery after injury show long-term impairment. **A:** In situ hybridization for *mpeg1* confirms the absence of macrophages before and after injury in the *irf8* mutant compared to controls. Arrows indicate labeling around the injury site and brackets indicate the ventral area of the larvae where the macrophages can be found in the circulation. Note that blackish colour is due to melanocytes. **B:** Quantification of the proportion of larvae with axonal bridging (anti-acetylated Tubulin) shows that at 1 dpl axonal bridging is unimpaired in *irf8* mutants, whereas at 2 dpl *irf8* mutants fail to show full regrowth and even by 5 dpl the proportion of *irf8* larvae with a bridged lesion site is still lower than in wildtype controls (*Fisher’s exact test*: ***P<0.001, n.s. indicated no significance). **C:** *Irf8* mutants never fully recover touch-evoked swimming in the observation period, whereas wildtype control animals do. Representative swim tracks are displayed. Note that unlesioned *irf8* larvae show swimming distances that are comparable to those in wildtype controls (*Two-way ANOVA*: F _15, 1372_ = 11.42, P<0.001; unles. = unlesioned, les. = lesioned) All lesions are done at 3 dpf. Lateral views of the injury site are shown; rostral is left. Arrows indicate axonal bridging. *Scale bars*: 200μm in A and 50μm in C. Error bars indicate SEM.

Next, we determined axonal and functional regeneration in *irf8* mutants compared to wildtype animals. In the wildtype situation, 44% of the animals showed continuous axon labelling across a spinal lesion site at 1 dpl. In *irf8* mutants the percentage was not different from controls (43%). At 2 dpl, however, axonal continuity was observed in 80% of wildtype animals but only in 41% of *irf8* mutants. (Fig. 2B). In order to see whether mutants would eventually recover, we assessed axonal regeneration at 5 dpl – 2.5 times as long as wildtype animals need for full axon bridging. Compared to 2 dpl, we found a slight increase in the proportion of larvae with bridged lesion sites (55% vs 41%), but regenerative success was still strongly reduced compared to wildtype controls (55% vs. 87% Fig. 2C).

Analysing touch-evoked swimming capacity, we found that wildtype animals swam distances that were comparable to those of unlesioned controls at 2 dpl, as previously described ^1^. In contrast, recovery in *irf8* larvae plateaued at 2 dpl and did not reach swim distances of unlesioned animals to at least 5 dpl (Fig. 2C). This indicates that in the absence of macrophages and microglia in *irf8* mutants, initial axon regeneration is unaffected, but full axon regrowth and functional recovery after spinal cord injury is long-term impaired.

To determine the relative importance of microglia for regeneration, we analysed *csf1ra/b* double-mutants (see Material and Methods) in which the function of colony-stimulating factor 1 receptor (Csf1r) is compromised. Csf1r is needed for microglia differentiation ^17^, but not for many macrophage functions ^18^. After injury in *csf1ra/b* mutants, we observed a strong reduction in the number of microglial cells (to 17% of wildtype), but an increase in macrophage numbers (by 55% compared to wildtype) in the injury site (Fig. 3A,C). Interestingly, neutrophil numbers were also reduced (to 16% of wildtype) (Fig. 3B), perhaps due to feedback regulation from increased macrophage numbers. Whereas microglia cells were reduced in number in the entire fish, neutrophils were still present in the ventral trunk area. In these mutants, axon bridging was unimpaired (Fig. 3D). Hence, microglia and sustained presence of neutrophils are not necessary for axon regeneration. Combined with results from the *irf8* mutant, this supports that recruitment of peripheral macrophages is critical for successful spinal cord regeneration.

**Fig. 3:**
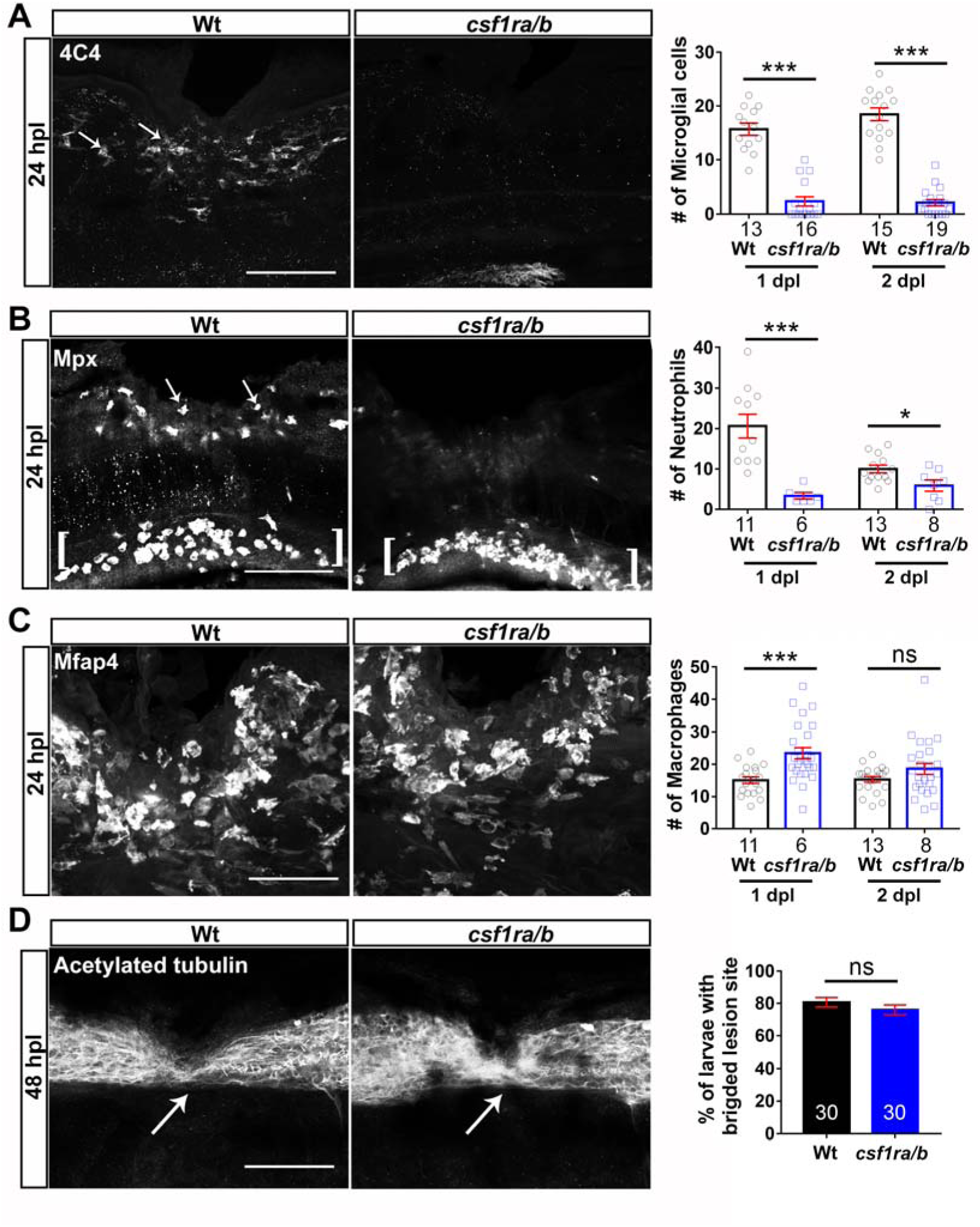
Absence of microglial cells and reduced neutrophil numbers do not affect axon bridging. **A:** Numbers of microglial cells (4C4^+^; arrows) in the injury site of the *csfr1a/b* mutants are much lower than in wildtype animals (*t-test*: ***P<0.001). **B:** Much fewer neutrophils (Mpx^+^) are found in the injury site (arrows) of *csfr1a/b* mutants than in wildtype animals (*t-test*: *P<0.05, ***P<0.001), but ventral neutrophils are still present (brackets). **C:** The number of macrophages (Mfap4^+^) is increased in the injury site in the mutants at 1dpl, but not at 2dpl (*t-test*: ***P<0.001, n.s. indicated no significance). **D:** Immunostaining against acetylated tubulin shows that axon bridging (arrows) is not affected in the mutants compared to wildtype animals at 2 dpl (*Fisher’s exact test*: n.s. indicated no significance). Lateral views of the injury site are shown; rostral is left. Wt = wildtype; Scale bars: 50μm in A, B, D; 25 μm in B. Error bars indicate SEM.

### Macrophages are not necessary for Col XII deposition

To determine the mechanism of action for macrophages, we tested for possible interactions with a previously reported regeneration-promoting mechanism, comprising Wnt-dependent deposition of Col XII in the lesion site by *pdgfrb*:GFP^+^ fibroblast-like cells ^1^. General inhibition of the immune response with dexamethasone did not inhibit appearance of *pdgfrb:*GFP^*+*^ fibroblast-like cells in the lesion site (Fig. S3A). By crossing a reporter line for Wnt pathway activity (*6xTCF*:dGFP) ^1^ into the *irf8* mutant, we found that activation of the pathway was unaltered in the mutant (Fig. S3E).

Similarly, expression of *col12a1a* and *col12a1b* mRNA in *irf8* mutants was indistinguishable from that in wildtype animals (Fig. S3B). Deposition of Col I protein and mRNA expression of 11 other ECM components were also not altered in the *irf8* mutant at 1 and 2 dpl (Fig. S3B,C). Moreover, immunolabelling against Tp63 showed that the injury site in the *irf8* mutants was completely covered again by basal keratinocytes, which are also producing Col XII^1^, by 2 dpl. This was similar to wildtype animals (Fig. S3D). Hence, macrophages do not overtly regulate Wnt-signalling, deposition of some crucial ECM components or keratinocyte coverage of the injury site during regeneration.

### Preventing increased cellular debris formation does not rescue lack of regenerative success in irf8 mutants

We hypothesized that macrophages were necessary to condition the environment through which axons grow, potentially by serving as a substrate for axon growth or by removing debris by phagocytosis - a major function of macrophages in peripheral nerve regeneration ^19,20^. In time-lapse movies of double transgenic animals in which neurons (*Xla.Tubb*:DsRed) and macrophages (*mpeg1*:GFP) were labelled (Fig. 4A and Movie S2) we observed axons crossing the spinal lesion site at the same time macrophages migrated in and out of the injury site. However, no obvious physical interactions between these cell types were observed, making it unlikely that macrophages acted as an axon growth substrate.

**Fig. 4:**
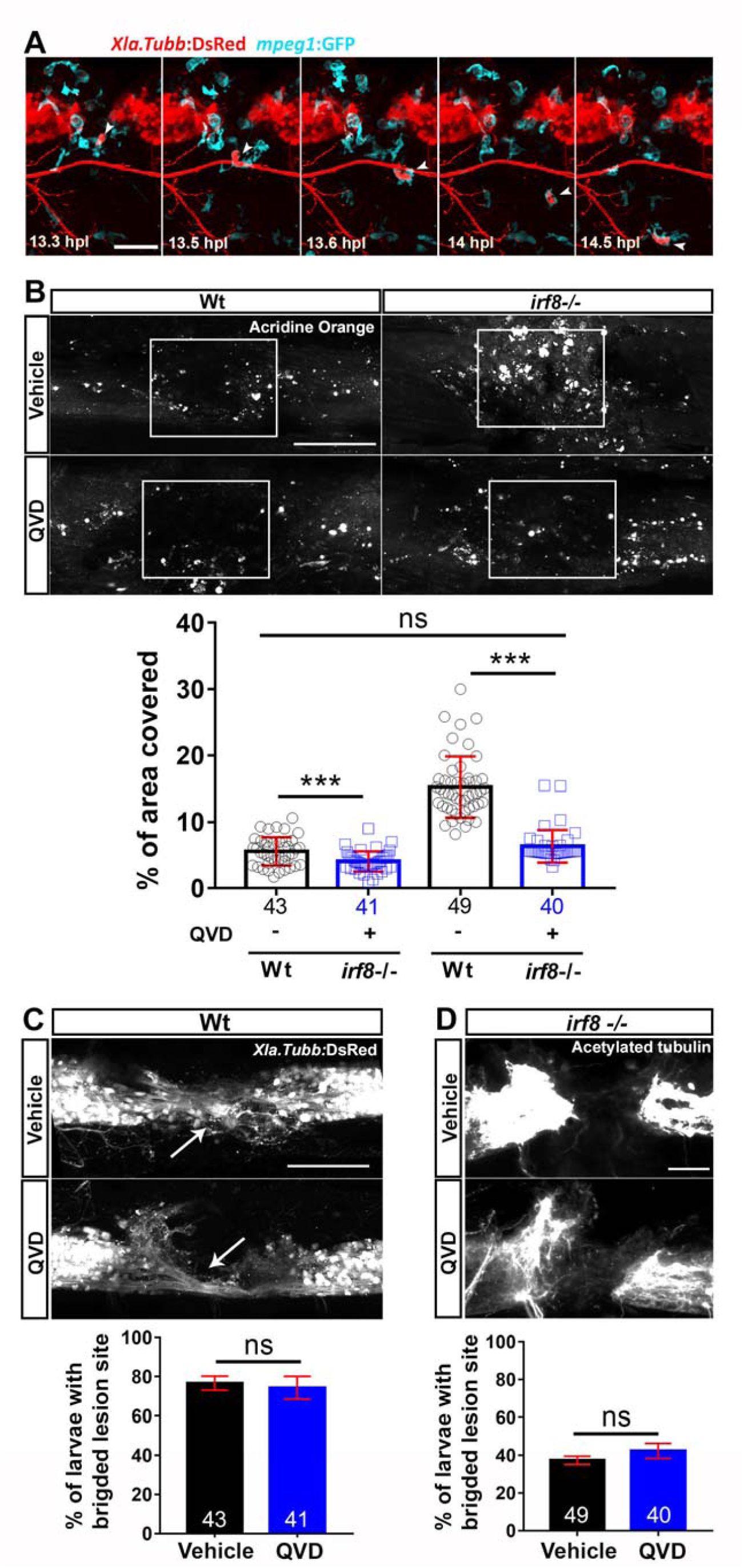
Debris removal by macrophages is not a major factor for axonal regeneration. **A:** Time-lapse video-microscopy shows that macrophages (cyan, *mpeg1*:EGFP) are in the injury site and remove neuronal debris (*Xla.Tubb*:DsRED). Arrowheads point out a macrophage with phagocytosed neuronal material. **B:** Acridine orange labelling indicates increased levels of debris in the *irf8* mutant, which can be reduced to wildtype levels with the pan-caspase inhibitor QVD at 2dpl (*Kruskal-Wallis with Dunn’s multiple comparisons post-test*: *** P<0.001, n.s. indicates no significance). **C,D:** Inhibition of cell death with QVD does not alter regenerative success in wildtype animals (C) or *irf8* mutants (D) at 2 dpl (*Fisher’s exact test*: n.s. indicates no significance). Rectangles indicate regions of quantification and arrows indicate axonal bridges. Lateral views of the injury site are shown; rostral is left. *Scale bars:* 100μm **A**, 50μm **B**, **C**, **D.** Error bars indicate SEM.

We frequently observed macrophages ingesting neuronal material and transporting that away from the injury site in time-lapse movies (Fig. 4A and Movie S2). In agreement with this observation, TUNEL labelling indicated strongly increased levels of dead or dying cells in the late (48 hpl), but not the early (24 hpl) phase of axonal regeneration in the injury site of *irf8* mutants (Fig. S4). This is consistent with the late manifestation of the axonal bridging phenotype in *irf8* mutants.

To determine the importance of debris for regeneration, we prevented debris accumulation in *irf8* larvae. We used the pan-caspase inhibitor QVD ^21^ to inhibit cell death. This treatment led to strongly reduced debris levels that were comparable to those seen in wildtype larvae at 2 dpl, as detected by acridine orange labelling (Fig. 4B). However, this treatment did not affect regenerative success in wildtype animals and it did not increase regenerative success in *irf8* mutants (control: 38% larvae with axon bridges; QVD: 40%) (Fig. 4C, D). This suggests that accumulation of debris is not the predominant cause for the failure of axons to regenerate across the injury site in *irf8* mutants.

### Pro-and anti-inflammatory phases are altered in irf8 mutants

To determine a possible role of cytokines in the regenerative failure of *irf8* mutants, we analysed relative levels of pro-and anti-inflammatory cytokines in the lesion site environment during regeneration in wildtype animals and *irf8* mutants by qRT-PCR. In wildtype animals, expression levels of pro-inflammatory cytokines *il-1β* and *tnf-*α were high during the initial phase of regeneration (> 25-fold for *il-1β*; >12-fold for *tnf-*α for approximately to 12 hpl) and expressed at lower levels during the late phase of regeneration (12-48 hpl), although still elevated compared to unlesioned age-matched controls (Fig. 5A,B). Anti-inflammatory cytokines, such as *tgf-β1a* and *tgf-β3* were expressed at low levels during initial regeneration, and upregulated during late regeneration (approximately 3-fold for *tgfb1and 2-fold for tgfβ3*), indicating a bi-phasic immune response within the 48-hour time frame of analysis (Fig. 5C,D).

**Fig. 5:**
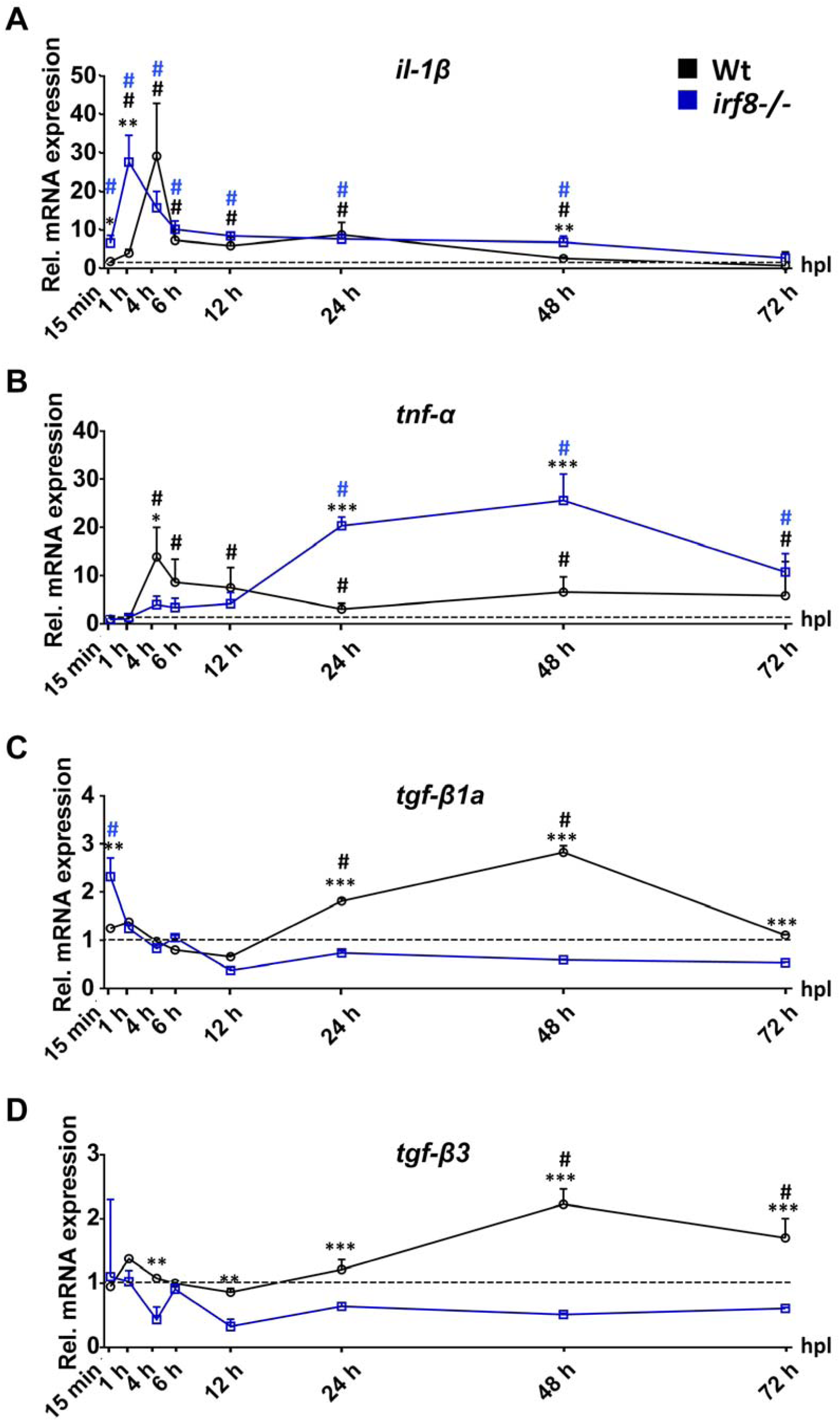
Inflammation is bi-phasic and dysregulated in *irf8* mutants. Absence of macrophages in the *irf8* mutant fish leads to increased *il-1β* and *tnf-*α mRNA levels during the late stage of inflammation (>12hpl). An early peak in *tnf-*α expression is missing in *irf8* mutants. Expression of anti-inflammatory cytokines, *tgfb-1a* and *tgfb3*, which peak during late regenerative phases in wildtype animals, is strongly reduced in *irf8* mutants (*t-tests*: *P<0.05, **P<0.01, ***P<0.001; wt = wildtype animals). # indicated statistical significance when compared to unlesioned animals. *Error bars* indicate SEM.

In *irf8* mutants, levels of pro-inflammatory cytokines remained high during the late phase of regeneration (Fig. 5A,B) and anti-inflammatory cytokines were not upregulated (Fig. 5C,D), resulting in a sustained pro-inflammatory environment in *irf8* mutants. Hence, the immune response is bi-phasic with an initial pro-inflammatory phase, followed by an anti-inflammatory phase in wildtype animals. In the absence of macrophages in *irf8* mutants, animals fail to switch to an anti-inflammatory state.

### TNF-α promotes axonal regeneration

As sustained inflammation is thought to be detrimental to spinal cord regeneration in mammals, we aimed to determine whether increased levels pf pro-inflammatory cytokines might contribute to impaired axon growth in *irf8* mutants. First, we inhibited TNF-α signalling with Pomalidomide, a pharmacological inhibitor of TNF-α release ^22^. However, this had no effect on axon regrowth in *irf8* mutants. In contrast, in wildtype animals Pomalidomide strongly inhibited axon bridging at 1 dpl (control: 62% bridging; Pomalidomide: 36%) and 2 dpl (control: 75% bridging; Pomalidomide: 45%) (Fig. 6A). To confirm the effect of TNF-α inhibition, we targeted *tnf-*α by using CRISPR manipulation with a gene–specific guideRNA (gRNA). Acute injection of the gRNA efficiently mutated the gene as shown by restriction fragment length polymorphism (RFLP) analysis (Fig. 6B). Axon bridging was inhibited in wildtype animals in a way that was comparable to drug treatment (1 dpl: control: 51% bridging; gRNA: 27%; 2 dpl: control: 88% bridging; gRNA: 40%; Fig. 6C) These results indicate that *tnf-α* upregulation is not a major cause of regenerative failure in *irf8* mutants, but is necessary for axon regeneration in wildtype animals.

**Fig. 6:**
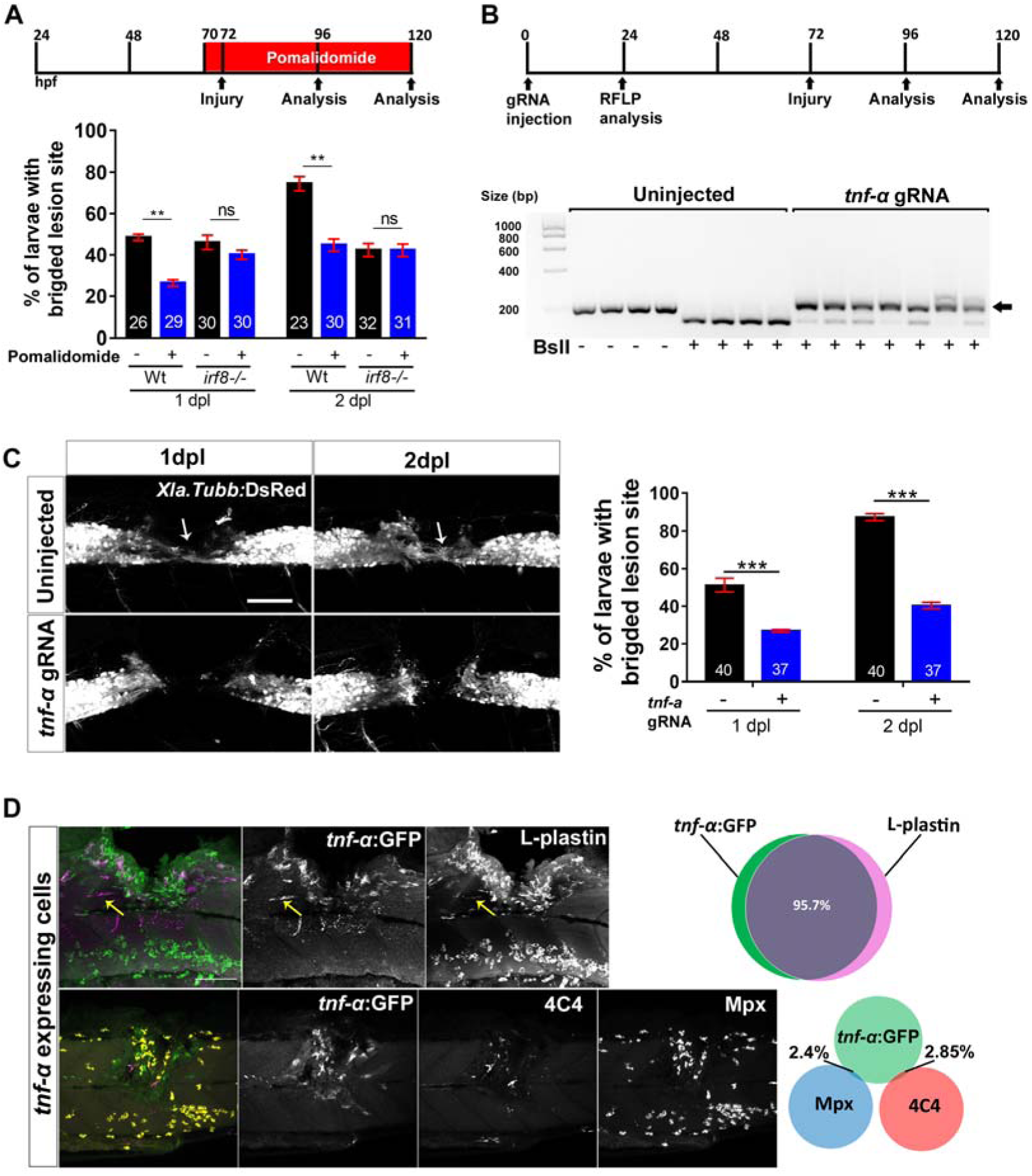
Tnf-α is essential for axonal regeneration. **A:** Tnf-α inhibition by Pomalidomide reduces the proportion of wildtype animals with axon bridging at 1 and 2 dpl. No effect is observed in *irf8* mutants (*Two-way ANOVA followed by Bonferroni post-test*: F _3, 16_ = 12.16, **P<0.01, n.s indicates no significance). **B:** CRISPR/Cas9-mediated disruption of *tnf-α* is effective as shown by RFLP analysis. This reveals efficient somatic mutation in the gRNA target site, indicated by resistance to restriction endonuclease digestion (arrow). **C:** Axonal bridging (arrow; *Xla.Tubb*:DsRed^+^) is impaired after disruption of the *tnf-α* gene. (*Fisher’s exact test*: **P<0.01). **D**: *tnf-α* is mostly expressed by peripheral macrophages in the lesion site of wildtype animals. Top row: *tnf-α*:GFP labelling occurs almost exclusively in L-plastin^+^ immune cells (L-plastin in green; *tnf-α*:GFP in purple; yellow arrows indicate a rare *tnf-* α:GFP^+^ microglial cell; 12 hpl) Bottom row: *tnf-α*:GFP (in green) rarely co-labels with microglial (4C4; in purple) or neutrophil (Mpx; in yellow) markers (24 hpl). Lateral views of the injury site are shown; rostral is left. *Scale bars:* 50μm. Error bars indicate SEM.

To determine which cells expressed *tnf-α* in wildtype animals, we used L-Plastin immunohistochemistry, labelling all immune cells, in *tnf-α*:GFP transgenic fish (Fig. 6D). This indicated that nearly all *tnf-α*:GFP^+^ cells co-labelled with the myeloid lineage marker L-Plastin (96%). Thus, expression of *tnf-α* occurred mainly in immune cells. Double-labelling *tnf-α*:GFP reporter fish with neutrophil (Mpx) and microglia (4C4) markers at 24 hpl, when axons were actively growing, indicated that 95% of *tnf-α*:GFP^+^ cells did not co-label with these markers (Fig. 6D). Hence, *tnf-α* was mostly expressed by peripheral macrophages (L-Plastin^+^, 4C4^-^, Mpx^-^). This supports the notion that macrophages promote regeneration by expressing *tnf-α*.

### Il-1β inhibits regeneration in irf8 mutants

To test whether sustained high levels of Il-1β were responsible for regenerative failure *irf8* mutants, we interfered with *il-1β* function in three different ways. Firstly, we inhibited caspase-1, which is necessary for activation of Il-1β. For this, we used the pharmacological inhibitor YVAD that is functional in zebrafish^23^ (Fig.7A-E). Secondly, we used an established morpholino to disrupt *il-1β* RNA splicing ^24^. Finally, we targeted *il-1β* with gRNA in a CRISPR approach (Fig. S5A.H).

To determine whether interfering with Il-1β function mitigated inflammation in *irf8* mutants, we quantified immune cells, expression of *il-1β* and tnf-α and dead cells. Indeed, after YVAD treatment we observed a reduction of neutrophil peak numbers (by 38% at 2 hpl; Fig. 7B), as well as reduced levels of *il-1β* and *tnf-α* expression (at 2 dpl; Fig. 7A) in *irf8* mutants. Moreover, the number of TUNEL^+^ cells was reduced at 2 dpl in the *irf8* mutant but was still much higher than in wildtype animals (Fig. 7C). In wildtype animals, incubation of lesioned larvae with YVAD reduced peak numbers of neutrophils (by 40% at 2 hpl) and macrophages (by 28% at 48 hpl), but no influence on low numbers of TUNEL^+^ cells at 2 dpl was observed. Hence, interfering with Il-1β function reduces inflammation in *irf8* mutants and wildtype animals

**Fig. 7:**
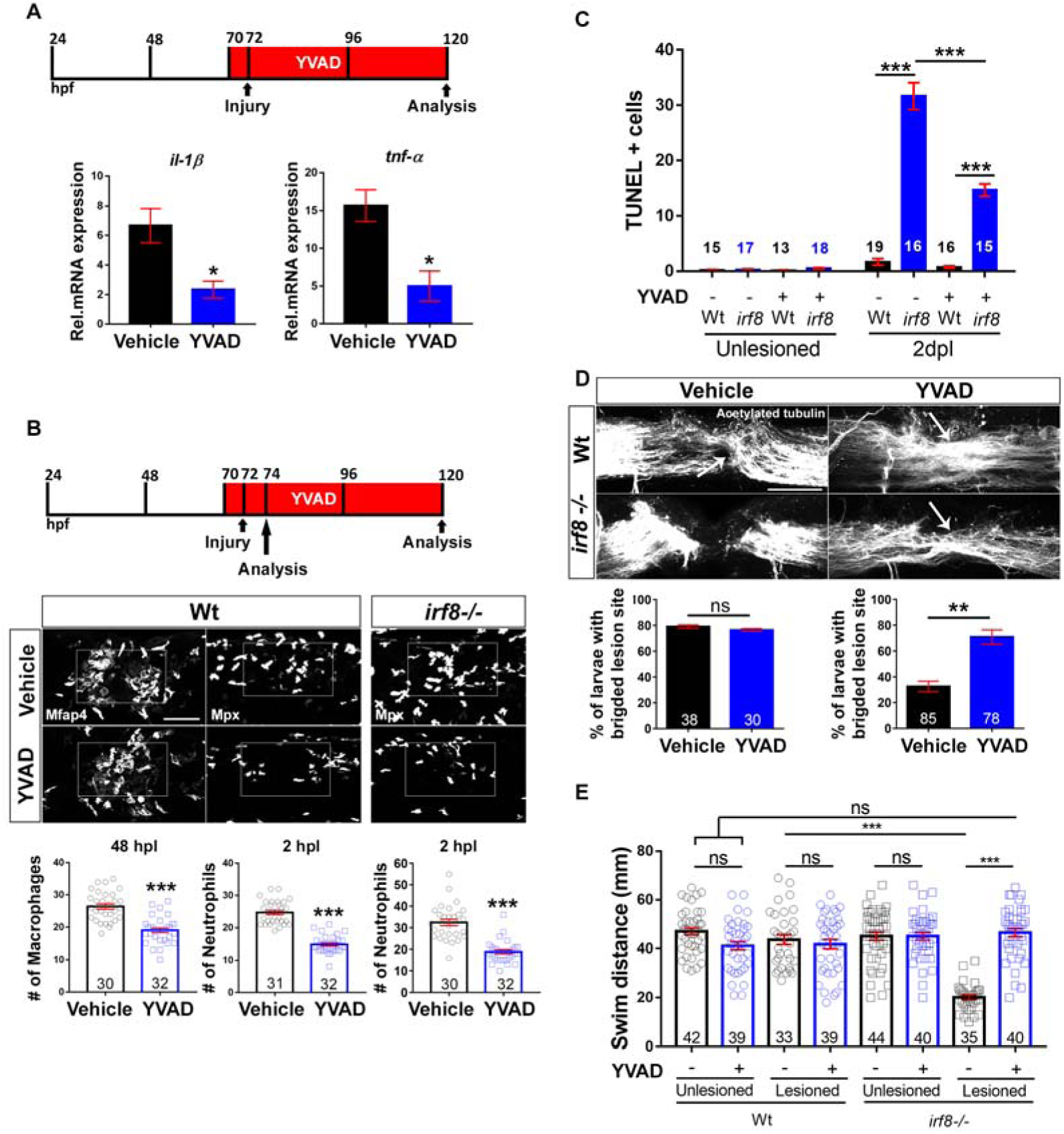
Inhibition of Il-1β function rescues axonal regeneration in *irf8* mutants. Lateral views of the injury site are shown; rostral is left. **A:** YVAD reduces expression levels of *il-1β* and *tnf-α* in *irf8* mutants (two-sample t-test: *P<0.05) at 2dpl. **B:** YVAD impairs migration of peripheral macrophages (Mfap4^+^) and neutrophils (Mpx^+^) in wildtype animals and *irf8* mutants (only neutrophils quantified, due to absence of macrophages) (t-tests: ***P<0.001). **C:** YVAD moderately reduces the number of TUNEL^+^ cells in the *irf8* mutants at 2dpl. (Two-Way ANOVA followed by Bonferroni multiple comparisons: F _3,_ 121 = 112.5, ***P<0.001). **D**: YVAD does not influence axonal regeneration in wildtype animals but rescues axonal bridging in *irf8* mutants (*Fisher’s exact test*: **P<0.01, n.s. indicated no significance) at 2 dpl. **E:** Inhibition of *il-1β* function restores functional recovery in *irf8* mutants. Impaired touch-evoked swimming in *irf8* mutants is rescues by YVAD treatment, to levels that are no longer different from lesioned and unlesioned wildtype animals at 2 dpl. YVAD has no influence on swimming in lesioned or unlesioned wildtype animals (*Two-way ANOVA followed by Bonferroni multiple comparisons*: F _1, 309_ = 35.229, ***P<0.0001, n.s. indicates no significance). Scale bar: 50μm for B, C. Error bars indicate SEM.

Axon bridging in wildtype animals was not affected by YVAD treatment at 2 dpl (control: 79% bridging; YVAD: 78% bridging) (Fig. 7D), indicating that high levels of Il-1β were not necessary for axonal regeneration. In contrast, in YVAD-treated *irf8* mutants, we observed a remarkable rescue of axon bridging at 2 dpl (con: 38% crossing; YVAD: 69% crossing) (Fig. 7D).

Injecting a well-established ^24^ morpholino targeting *il-1β* into *irf8* mutants at the one-cell-stage inhibited *il-1β* splicing (Fig. S5A,D). Consistent with pharmacological data, morpholino-injected animals showed a rescue of the axon bridging phenotype at 2 dpl (con: 40% crossing; YVAD: 60% bridging) (Fig. S5B,C).

Finally, injecting a gRNA targeting *il-1β* at the one-cell stage led to somatic mutation in the target site of *il-1β*, indicated by RFLP analysis (Fig. S5E,H). We observed a strong rescue of axonal bridging in gRNA-injected. lesioned *irf8* mutants (con: 40% bridging; acute *il-1β* gRNA: 70% bridging) (Fig. S5F,G) by this treatment. Hence, three independent manipulations show that excessive *il-1β* levels in *irf8* mutants are a key reason for impaired axonal regeneration.

### Il-1β promotes axonal regeneration during the early phase of regeneration

To determine whether roles of *Il-1β* and general inflammation differed for different phases of the inflammation, we separately analysed early (0-1 dpl; Fig. S6A) and late (1-2 dpl; Fig. S6B) regeneration by drug incubation. During the early phase, YVAD treatment led to a weak inhibition of axon regeneration in both wildtype (con: 58% crossing; YVAD: 41% crossing) and *irf8* mutants (con: 41% crossing; YVAD: 36% crossing). Similarly, dexamethasone treatment inhibited axon regeneration in both wildtype (control: 57.5%; Dex: 36.6%) and *irf8* (control: 44.6%; Dex: 34%). Interestingly, while LPS promoted regeneration in the early phase in wildtype animals (control: 53.2%; LPS: 68.7%), it was detrimental in *irf8* mutants (control: 39%; LPS: 25.5%), perhaps because baseline inflammation was already high in the mutant.

During late regeneration, only dexamethasone had an inhibitory effect in wildtype animals (from 82.1% bridging to 64.4%). YVAD had no effect in wildtype animals during late regeneration, when *il-1β* was already down-regulated (control: 81% crossing; YVAD: 78% crossing), but a strong rescue effect of YVAD was observed in the *irf8* mutant (con: 37% crossing; YVAD: 68% crossing). This rescue effect was comparable to that observed when Il-1β was suppressed for the entire 48 hours (cf. Fig. 7C). LPS had no effect in wildtype or mutants during late regeneration. Taken together, these observations support that early inflammation and *il-1β* upregulation promote regeneration but *il-1β* must be down-regulated at later phases of axonal regeneration.

### Reduction of Il-1β levels fully rescues swimming deficits in irf8 mutants

To determine whether rescued axon regeneration after inhibition of *il-1β* also led to functionally meaningful regeneration in *irf8* mutants, we performed swimming tests using YVAD to inhibit Il-1β. The treatment had no effect on the swimming distances in unlesioned wildtype and *irf8* mutants and did not affect recovery in lesioned wildtype animals (Fig. 7E), in agreement with a lack of an overall effect on axonal regrowth (cf. Fig. 7B). In contrast, treatment with YVAD rescued the behavioural impairment in the *irf8* mutants to levels that were indistinguishable from wildtype lesioned or unlesioned animals (Fig. 7E). These observations indicate that experimental reduction of *Il-1β* alone is sufficient to restore most axonal regeneration and full recovery in touch-evoked swimming in the absence of macrophages.

### Neutrophils are a major source of il-1β and inhibit regeneration

To understand how *il-1β* was regulated, we determined the cellular source of Il-1β in wildtype and *irf8* mutants. Detection of *il-1β* mRNA by qRT-PCR and in situ hybridisation confirmed increased levels at 2 dpl, but not 1 dpl in the *irf8* mutant (Fig. 8A,B). Using Il-1β immunohistochemistry and a transgenic reporter line (*il-1β*:GFP), we found that many neutrophils and keratinocytes were positive for Il-1β in wildtype and *irf8* mutants at 1 dpl. Macrophages (Mfap4^+^) were frequently positive for Il-1β in wildtype animals, whereas neurons (HuC/D^+^) were rarely co-labelled (Fig. S7A,B). Increased numbers of Il-1β^+^ neutrophils (by 97%) and keratinocytes (by 58%) were observed the *irf8* mutant, consistent with the higher pro-inflammatory status of *irf8* mutants (Fig. 8C,D). Importantly, the proportion of neutrophils that were Il-1β^+^ was also increased from 28% in wildtype animals to 49% in the *irf8* mutant. This demonstrates that neutrophils are more likely to express *il-1β* in the absence of macrophages.

**Fig. 8:**
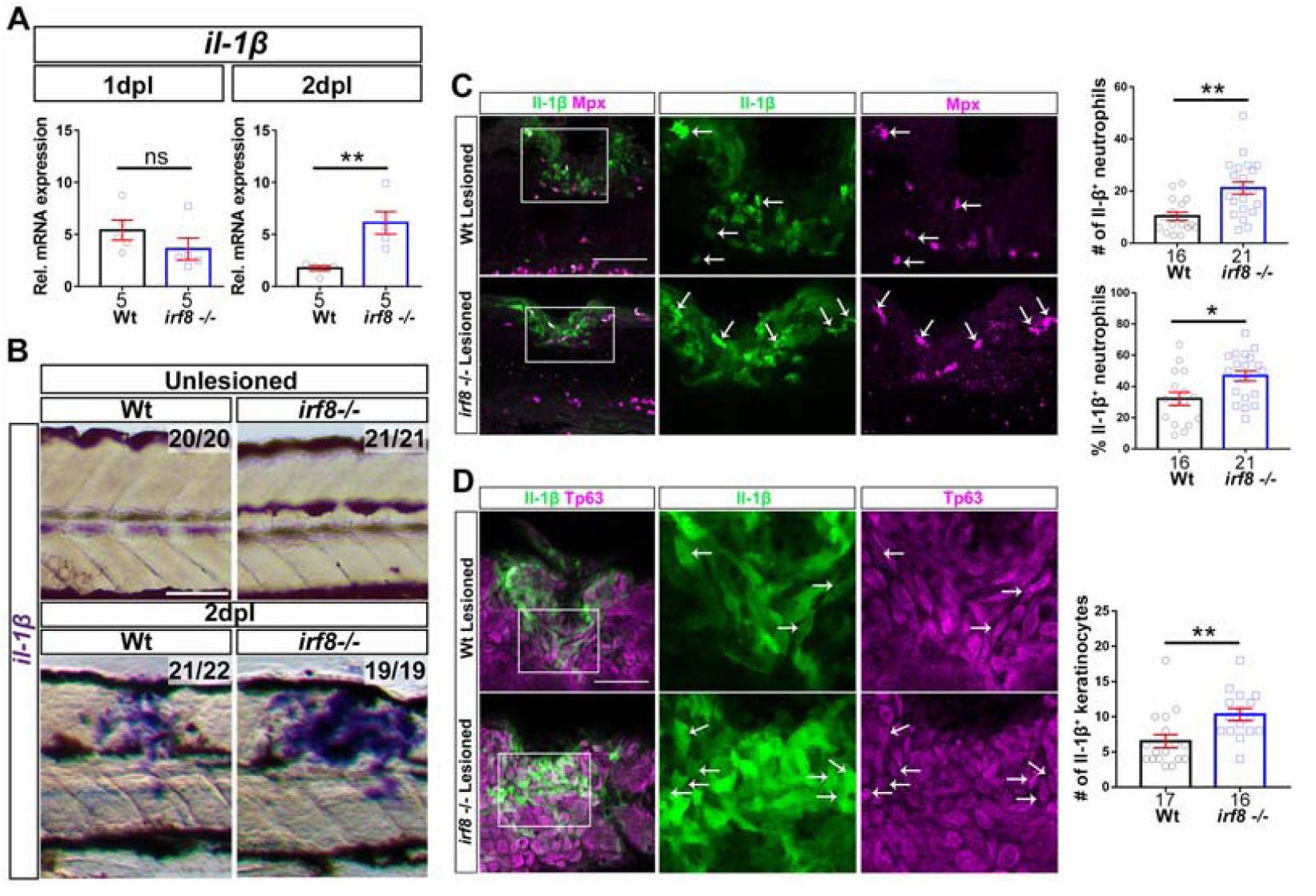
Levels of *il-1β* expression are increased in the injury site of *irf8* mutants at 2dpl. Lateral views of the injury site are shown; rostral is left**. A:** At 1 dpl, expression levels of *il-1β* are comparable between *irf8* mutants and wildtype (Wt) animals but are higher in the mutant at 2 dpl in qRT-PCR (*t-test*: **P<0.01, n.s. indicated no significance). **B:** In situ hybridization confirms increased expression of *il-1β* mRNA at 2 dpl. **C,D:** In the injury site, the number and proportion of neutrophils (Mpx^+^) that are Il-1β immuno-positive (arrows) are increased in *irf8* mutants compared to wildtype animals (C). The number of keratinocytes (Tp63^+^) that are Il-1β immuno-positive is increased in irf8 mutants (D). Single optical sections are shown; boxed areas are shown in higher magnifications; 1 dpl; *t-test*: *P<0.05, **P<0.01). Lateral views of the injury site are shown; rostral is left. Scale bars: 100μm in B,C,D and 50 μm for higher magnification areas. Error bars indicate SEM.

Increased numbers of Il-1β^+^ neutrophils in *irf8* mutants could, at least in part, be due to higher overall numbers of neutrophils in the injury site. We found a peak of neutrophil numbers in the lesion site of *irf8* mutants at 2 hpl, as in wildtype animals (Fig. 9A). However, the number of neutrophils was 27 % higher than in wildtype, potentially due to the higher abundance of this cell type in the *irf8* mutant ^16^. While neutrophil numbers declined over time in wildtype and *irf8* mutants, they did so more slowly in *irf8* mutants. At 24 hpl, twice as many neutrophils remained at the lesion site and at 48 hpl, three times the number of neutrophils as in wildtypes remained in the mutant. These observations support that macrophages have role in controlling cytokine expression status and number of neutrophils at the injury site ^25^. Hence, *irf8* mutants show prolonged presence of Il-1β^+^ neutrophils in the spinal lesion site.

**Fig. 9:**
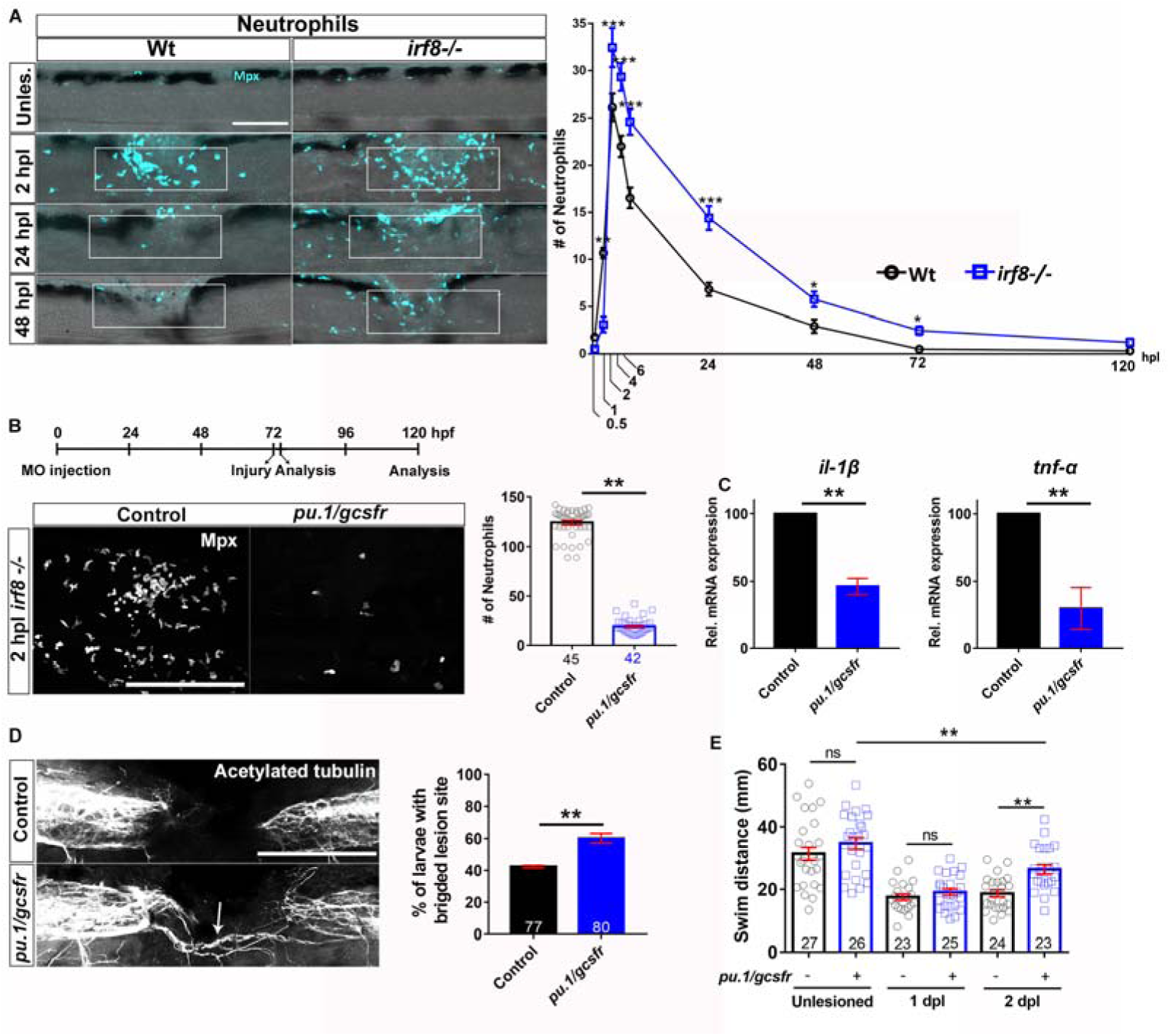
Preventing neutrophil formation partially rescues functional spinal cord regeneration in the *irf8* mutant. Lateral views of the injury site are shown; rostral is left. **A**: In *irf8* mutants, higher peak numbers of neutrophils (Mpx^+^) at 2 hpl and slower clearance over the course of regeneration are observed (*Two-Way ANOVA followed by Bonferroni multiple comparisons*: F _8, 427_ = 13.19 *P<0.05, **P<0.01, ***P<0.001). Note that wildtype data are the same as shown in Fig. 1A, as counts in *irf8* mutants and wildtype animals were done in the same experiments. **B:** Combination treatment with *pu.1* and *gcsfr* morpholinos efficiently prevents neutrophil accumulation in the lesion site (Mann-Whitney U-test: ***P<0.001). **C**: In *pu.1/gcsfr* morpholino injected *irf8* mutant fish, levels of *il-1β* and *tnf-α* mRNA expression are reduced at 2 dpl, as shown by qRT-PCR (t-test: ***P<0.001). **D,E:** In *pu.1/gcsfr* morpholino injected *irf8* mutant fish, axonal bridging (D; *Fisher’s exact test*: **P<0.01) and behavioural recovery (E; *One-Way ANOVA followed by Bonferroni multiple comparisons*: F _5, 142_ = 23.21, **P<0.01, n.s. indicates no significance) are partially rescued. *Error bars* indicate SEM. *Scale bars:* 100μm.

In order to determine the relative importance of the neutrophils for regenerative failure in *irf8* mutants, we reduced their numbers using *pu.1*/*gcsfr* morpholino treatment. Because *irf8* mutants did not possess macrophages or microglia, this treatment specifically inhibited neutrophil formation, essentially creating a fish without a local immune response. This treatment strongly reduced numbers of neutrophils by 84.6% at 2 hpl in the injury site, when the neutrophil reaction peaked in untreated *irf8* mutants (Fig. 9B). Preventing formation of neutrophils in *irf8* mutants also reduced *il-1β* mRNA levels by 54% and *tnf-α* mRNA levels by 70% at 2 dpl (Fig. 9C). This indicates that neutrophils are a major source of excess *il-1β* in *irf8* mutants.

Remarkably, axon bridging was partially rescued in these neutrophil-depleted mutants (control: 43%; *pu.1*/*gcsfr* morpholino: 60%; Fig. 9D). Moreover, recovery of swimming behaviour was not altered at 1 dpl (when it is similar between wild type animals and *irf8* mutants), but was partially rescued at 2 dpl (Fig. 9E). This shows that in the absence of macrophages, the prolonged presence of Il-1β^+^ neutrophils is detrimental to regeneration.

## DISCUSSION

In this study we identify a biphasic role of the innate immune response for axonal bridging of the non-neural lesion site in larval zebrafish. Initial inflammation and the presence of Il-1β promote axon bridging, whereas during late regeneration, Il-1β levels need to be tightly controlled by peripheral macrophages. Inhibiting Il-1β was sufficient to compensate for the absence of macrophages, underscoring the central role of this cytokine. Macrophage-derived Tnf-α promoted regeneration (schematically illustrated in Fig. 10). This indicates important and highly dynamic functions of the immune system for successful spinal cord regeneration.

**Fig. 10:**
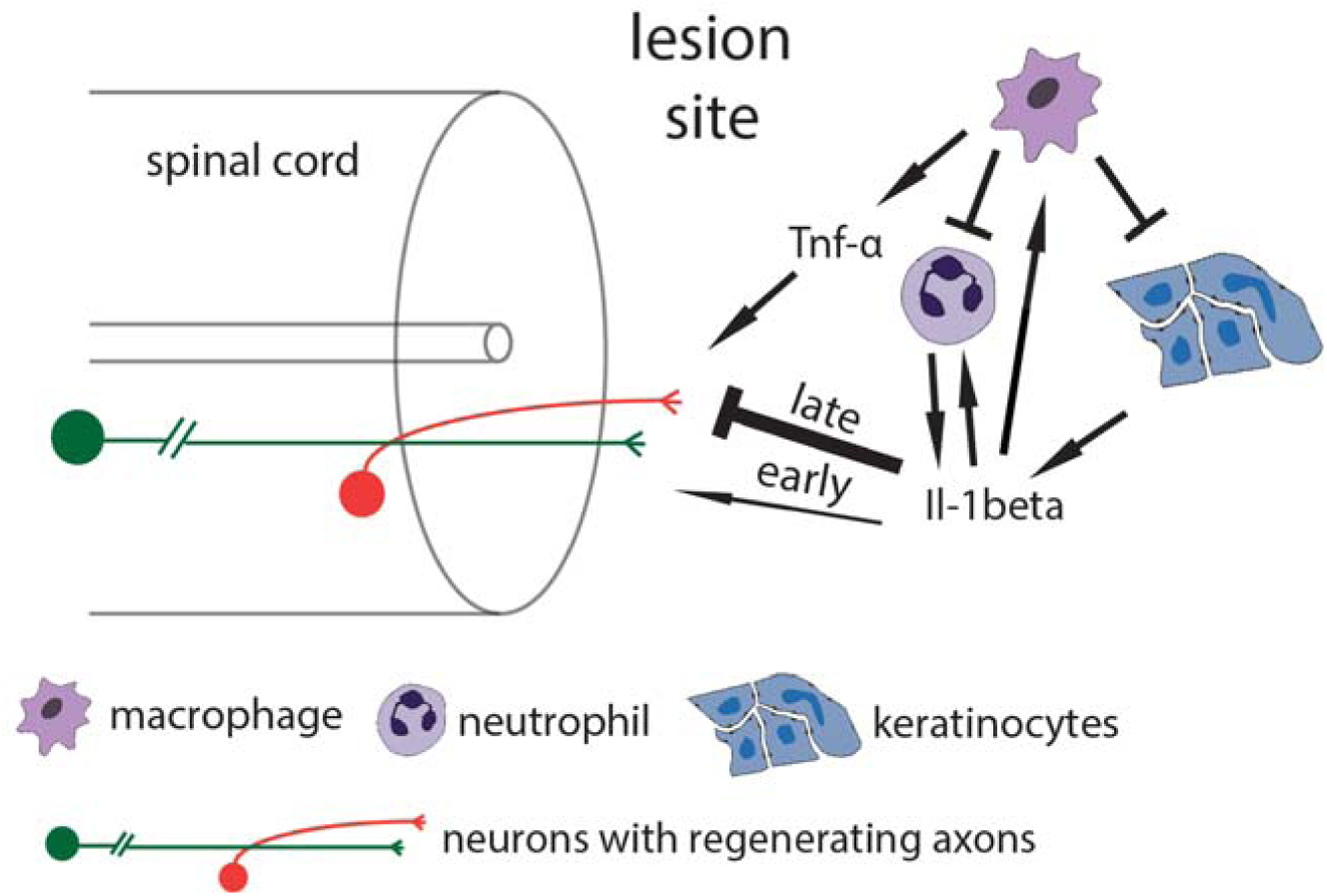
Working model of the influence of the innate immune system on axon regrowth. Spinal cord injury triggers neutrophil invasion of the injury site and initiation of the inflammatory response, including *il-1β* production. This initially promotes regeneration but is later strongly inhibitory. Il-1β provides positive feedback for neutrophil and macrophage presence. Macrophages invade the lesion site and down-regulate levels of *il-1β* in neutrophils and keratinocytes, while promoting regeneration by releasing Tnf-α. This leads to successful axonal regeneration and functional recovery.

The function of the immune system in spinal cord regeneration changes dramatically over time. Within the first hours after injury, neutrophils and the pro-inflammatory cytokines *il-1β* and *tnf-α* dominate the injury site. At the early time points, inflammation has beneficial effects on axonal growth. This is indicated by reducing effects on axon bridging of early Il-1β inhibition in wildtype and *irf8* mutant fish and the promoting effect of LPS in wildtype animals. However, as of about 12 hpl, macrophages and anti-inflammatory cytokines are present in the lesion site and during this late phase of regeneration, Il1-β mediated inflammation in macrophage-less mutants strongly inhibits regeneration.

How does Il-1β inhibit spinal cord regeneration? It could act directly on axons, as it has been reported for sensory axons in mice ^26^. However, it is also possible that high Il-1β levels condition the environment to be inhibitory to axon regrowth. Excess Il-1β increases numbers of neutrophils, levels of *il-1β* and *tnf-α* expression, as well as cell death in the lesion environment. This is indicated by reductions in these parameters by blocking Il-1β signalling in *irf8* mutants. However, several ECM mutant, such that potentially detrimental down-stream effects of increased *il-1β* signalling in the environment remain to be determined. Similar to our observations, an Il-1β deficient mouse showed slightly increased axon regrowth after spinal injury ^27^.

Axonal regeneration is promoted by Tnf-α. Even though *il-1β* and *tnf-α* were similarly upregulated in the *irf8* mutant, only reducing Il-1β levels, but not Tnf-α levels rescued the mutant. Conversely, in wildtype animals, lack of Il-1β function had only a relatively small inhibitory effect on early regeneration, whereas Tnf-α was indispensable for axon regrowth. In fin regeneration, Tnf-α has an important promoting function for blastema formation ^28^. This suggests that Tnf-α may be involved in remodelling repair cells in the lesion site after spinal injury, which then creates an axon growth-promoting environment. The role of Tnf-α for axon regeneration in mammals is not clear and may be either promoting ^29,30^ or inhibiting ^31^.

Neutrophils, expressing *il-1β*, are major mediators of the inhibitory immune response together with keratinocytes in the absence of macrophages. This is indicated by reduced *il-1β* expression and the rescue of axon regrowth and swimming function when neutrophils were prevented from being formed in *irf8* mutants. However, the rescue was only partial. This is likely due to a combination of the absence of the regeneration-promoting function of the early immune response and presence of keratinocytes that may still express *il-1β* in neutrophil-depleted *irf8* mutants at later time points. In mammals, it has been proposed that neutrophils cause secondary cell death ^32^ and that depleting neutrophils leads to more favourable injury outcomes ^6^, similar to our observations.

Macrophages control inflammation, as their absence in *irf8* mutants leads to abnormally high expression levels of pro-inflammatory cytokines *il-1β* and *tnf-α*. This is similar to observations in fin regeneration ^33^. In the absence of macrophages, positive feedback regulation of *il-1β* takes place, as indicated by more *il-1β* positive neutrophils and keratinocytes in the *irf8* mutants and reduced *il-1β* mRNA levels when Il-1β function was inhibited. Moreover, a higher proportion of neutrophils were Il-1β^+^ in *irf8* mutants, showing that without macrophages, neutrophils have a more pro-inflammatory phenotype. Strongly reduced levels of anti-inflammatory cytokines *tgfβ1a* and *tgfβ3* in the *irf8* mutant suggest that these cytokines could be derived from macrophages and be responsible for reducing pro-inflammatory phenotypes in wildtype animals.

Macrophages do not promote regeneration primarily by preventing cell death or removing debris. Debris removal by phagocytosis is one of the major functions of macrophages and we observed this by time-lapse imaging in the present system after injury. In the *irf8* mutant, numbers of dead or dying cells were increased. Even under regeneration-permissive conditions, when *il-1β* was inhibited, levels of debris were still higher than in wildtype controls. Hence, after inhibition of *il-1β* in the *irf8* mutants, axons grew through a debris-filled lesion site. Moreover, preventing debris formation in *irf8* mutants using a pan-caspase inhibitor did not rescue regeneration. These data show that regenerative success does not correlate with debris abundance. Interestingly, in fin regeneration, lack of macrophages also leads to increased cell death. As this leads to death of tissue progenitor cells, fin regeneration is inhibited ^33^. In mammalian spinal injury, cellular debris, especially myelin debris, is seen as inhibitory to regeneration ^34,35^.

Are macrophages sufficient for axonal regrowth? Unimpaired axon regrowth in the *csfr1a/b* mutant, in which microglial cells and neutrophils are drastically reduced in number indicates that these cells may be dispensable for regeneration. However, the increase in peripheral macrophages in this mutant could have compensated for a possible contribution of microglia to axon regrowth. Moreover, neutrophils could be important immediately after injury and have systemic effects in the *csfr1a/b* mutant. It will be important to determine which signals from macrophages may fully support spinal cord regeneration through the non-neural lesion site environment.

Endothelial cells and myelinating cells are unlikely to be major mediators of regeneration in larval spinal cord regeneration. Endothelial cells from injured blood vessels were slow to reform blood vessels and were rarely invading the lesion site. In contrast, in mammals endothelial cells accumulate in the injury site, where they may have anti-inflammatory functions ^36^. Myelinating cells bridged the lesion site, but were not abundant and only did so, when axons had already formed bridges across the lesion site in most animals. In mammals, transplanted myelinating cells, such as olfactory ensheathing cells and Schwann cells have been shown to improve recovery after spinal injury ^37,38^.

Timing of the immune response is crucial for regenerative success after spinal lesion. Macrophages in mammals ^39^ and zebrafish ^40^ display pro-and anti-inflammatory phenotypes and the anti-inflammatory phenotypes are seen as beneficial for regeneration. We show that inflammation is rapidly down-regulated in zebrafish concurrent with the upregulation of anti-inflammatory cytokines, which does not readily occur in mammals ^4^.

We have established a functionally regenerating in vivo system in which complex interactions of immune cells and a spinal injury site can be rapidly observed and manipulated. This allows fundamental insight into the role of immune cells in successful axon regeneration that may ultimately inform non-regenerating systems.

We here use this system to demonstrate a pivotal role of macrophages in controlling Il-1β mediated inflammation and thereby promoting functional spinal cord regeneration.

## MATERIAL AND METHODS

### Animals

All zebrafish lines were kept and raised under standard conditions ^41^ and all experiments were approved by the British Home Office (project license no.: 70/8805). Regeneration proceeds within 48 hours of the lesion, therefore most analyses of axonal regrowth, cellular repair, and behavioural recovery can be performed before the fish are protected under the A(SP)A 1986, reducing the number of animals used in regeneration studies following the principles of the 3Rs. Approximately 10000 larvae of either sex were used for this study, of which 8% were over 5 dpf.

The following lines were used: WIK wild type zebrafish, Tg(*Xla.Tubb:*DsRed)^zf14826^, abbreviated as *Xla.Tubb:*DsRed ^42^; Tg(*mpeg1:*EGFP)^gl22^, abbreviated as *mpeg1*:GFP ^43^, and Tg(*mpx:*GFP)^uwm1^, abbreviated as *mpx:*GFP ^44^, Tg(*fli1:*EGFP)^y1^, abbreviated as *fli1*:GFP ^45^, *irf8*^st95/st95^ ^16^, TgBAC(*pdgfrb:*Gal4FF)^ncv24^;Tg(*UAS*:GFP), abbreviated as *pdgfrb*:GFP ^46^,Tg(*6xTCF/LefminiP*:2dGFP), abbreviated as *6xTCF*:dGFP ^47^, Tg(*claudin k*:Gal4)^ue101^; Tg(14x*UAS*:GFP) abbreviated as *cldnK*:GFP ^48^, Tg(*tnfa*:eGFP-F)^sa43296^, abbreviated as *tnf-α*:GFP ^40^ and Tg(*il-1β*:eGFP)^sh445^ ^49^, abbreviated as *il-1β*:GFP.

### Generation of new mutant and transgenic lines

Zebrafish deficient for *csf1ra* and *csf1rb* were generated by incrossing *csf1ra*^*j4e1*/j4e1^ with a V614M substitution in the first kinase domain ^50^ to a deletion allele of *csf1rb* generated by TALEN. The TALEN arms targeted exon 3 of the *csf1rb* gene, resulting in a 4 bp deletion and premature stop codon (Oosterhof et al., in preparation). We used incrosses of *csfr1a-/-; csfr1b*^*+*^*/-* for our analyses. We excluded larvae from microglial cells (4C4 co-labeling) in the head that were similar to those found in wildtype animals. All other larvae had either very few or no microglial cells in their heads. We refer to the double mutants as *csfr1a/b* mutants throughout the manuscript.

### Drug treatment

Dexamethasone (Dex) (Sigma) was dissolved in DMSO to a stock concentration of 5mM. The working concentration was 10 μM prepared by dilution from stock solution in fish water. Ac-YVAD-cmk (YVAD) (Sigma) was dissolved in DMSO to a stock concentration of 10 mM. The working concentration was 50 μM prepared by dilution from the stock solution in fish water. Q-VD-OPh (Sigma), abbreviated as QVD in the manuscript, was dissolved in DMSO to a stock concentration of 10 mM. The working concentration was 50 μM.

Lipopolysaccharides from Escherichia coli O55:B5 (LPS, Sigma) were dissolved in PBS to a stock concentration of 1 mg/ml. The working dilution was 50 μg/ml. Pomalidomide (Cayman Chemicals) was diluted in DMSO at a stock concentration of 10mg/ml. For the treatments, 6.9 μl of the stock where diluted in 1.5 ml of fish water. Larvae were pre-treated for 2 hours before the injury and were incubated for 24 and 48 hpl. Larvae were collected from the breeding tanks and were randomly divided into Petri dishes at a density of maximally 30 larvae per dish, but no formal randomization method was used. For the Dex experiment the larvae were treated from 6 hours post fertilization (hpf) until 5 dpf. For most drug treatments, larvae were incubated with the drug from 3 dpl until 5 dpl, if not indicated differently.

### Spinal cord lesions

Zebrafish larvae at 3 dpf were anesthetized in PBS containing 0.02% aminobenzoic-acid-ethyl methyl-ester (MS222, Sigma), as described ^51^. Larvae were transferred to an agarose-coated petri dish. Following removal of excess water, the larvae were placed in a lateral position, and the tip of a sharp 30 ½ G syringe needle was used to inflict a stab injury or a dorsal incision on the dorsal part of the trunk at the level of the 15^th^ myotome.

### Behavioural analysis

Behavioral analysis was performed as previously described^14^. Briefly, lesioned and unlesioned larvae were touched caudal to the lesion site using a glass capillary. The swim distance of their escape response was recorded for 15 secs after touch and analyzed using a Noldus behavior analysis setup (EthoVision version 7). Data given is the average distance travelled from triplicate measures per fish. Between repeated measures, the larvae were left to recover for 1 min. The observer was blinded to the treatment during the behavioral assay. The assay was performed on five independent clutches in order to assess the behavioral recovery in the *irf8* mutants and three independent clutches after the YVAD treatment.

### Quantitative RT-PCR

RNA was isolated from the injury sites of the larvae using the RNeasy Mini Kit (Qiagen). Forty larvae were used for each condition. cDNA used as template was created using the iScript™ cDNA Synthesis Kit (Bio-Rad). Standard RT-PCR was performed using SYBR Green Master Mix (Bio-Rad). qRT-PCR was performed at 58°C using Roche Light Cycler 96 and relative mRNA levels determined using the Roche Light Cycler 96 SW1 software. Samples were run in duplicates and expression levels were normalized to β-actin control. Primers were designed to span an exon exon junction using Primer-BLAST. Sequences are given in table 1. All experiments were carried out at least as biological triplicates.

**Table 1:** Antibody list

| **Antibody** | **Raised in** | **Source** | **Catalog #** | **Dilution** |
| --- | --- | --- | --- | --- |
| anti-GFP<sup>1</sup> | chicken | Abcam | AB13970 | 1:300 |
| Anti-4C4<sup>14</sup> | mouse | European Collection of Authenticated Cell Cultures (ECACC) | 7.4.C4 (ECACC 92092321) | 1:50 |
| Monoclonal Anti-Tubulin, Acetylated<sup>1</sup> | mouse | Sigma | T6793 | 1:300 |
| Anti-Tp63<sup>1</sup> | rabbit | Sigma | SAB2701838 | 1:300 |
| Anti-Col I<sup>1</sup> | rabbit | Abcam | ab23730 | 1:300 |
| Anti-HuC/HuD<sup>61</sup> | mouse | Invitrogen | A-21271 | 1:100 |
| Anti-Mpx<sup>62</sup> | rabbit | GeneTex | GTX128379 | 1:300 |
| Anti-Mfap4 | rabbit | GeneTex | GTX132692 | 1:300 |
| Anti-II-1β<sup>63</sup> | rabbit | Proteintech | 16806-1-AP | 1:200 |
| Anti-L-plastin<sup>14</sup> | rabbit | Yi Feng | - | 1:500 |
| AffiniPure Fab Fragment Donkey Anti-Rabbit IgG (H<sup>+</sup>L)<sup>52,64</sup> | donkey | Jackson ImmunoResearch | 711-007-003 | 40 μg/ml |
| Cy5 Anti-rabbit | donkey | Jackson ImmunoResearch | 711-175-152 | 1:300 |
| Cy<sup>TM</sup>3 Anti-Rabbit IgG (H<sup>+</sup>L) | donkey | Jackson ImmunoResearch | 711-165-152 | 1:300 |
| Cy<sup>TM</sup>3 Aff Anti-Mouse IgG (H<sup>+</sup>L) | donkey | Jackson ImmunoResearch | 715-165-150 | 1:300 |
| Alexa Fluor® 488 Anti-Mouse IgG(H<sup>+</sup>L) | donkey | Jackson ImmunoResearch | 715-545-150 | 1:300 |
| Cy<sup>TM</sup>5- Anti-Mouse IgG (H<sup>+</sup>L) | donkey | Jackson ImmunoResearch | 715-175-150 | 1:300 |

**Table 2:**
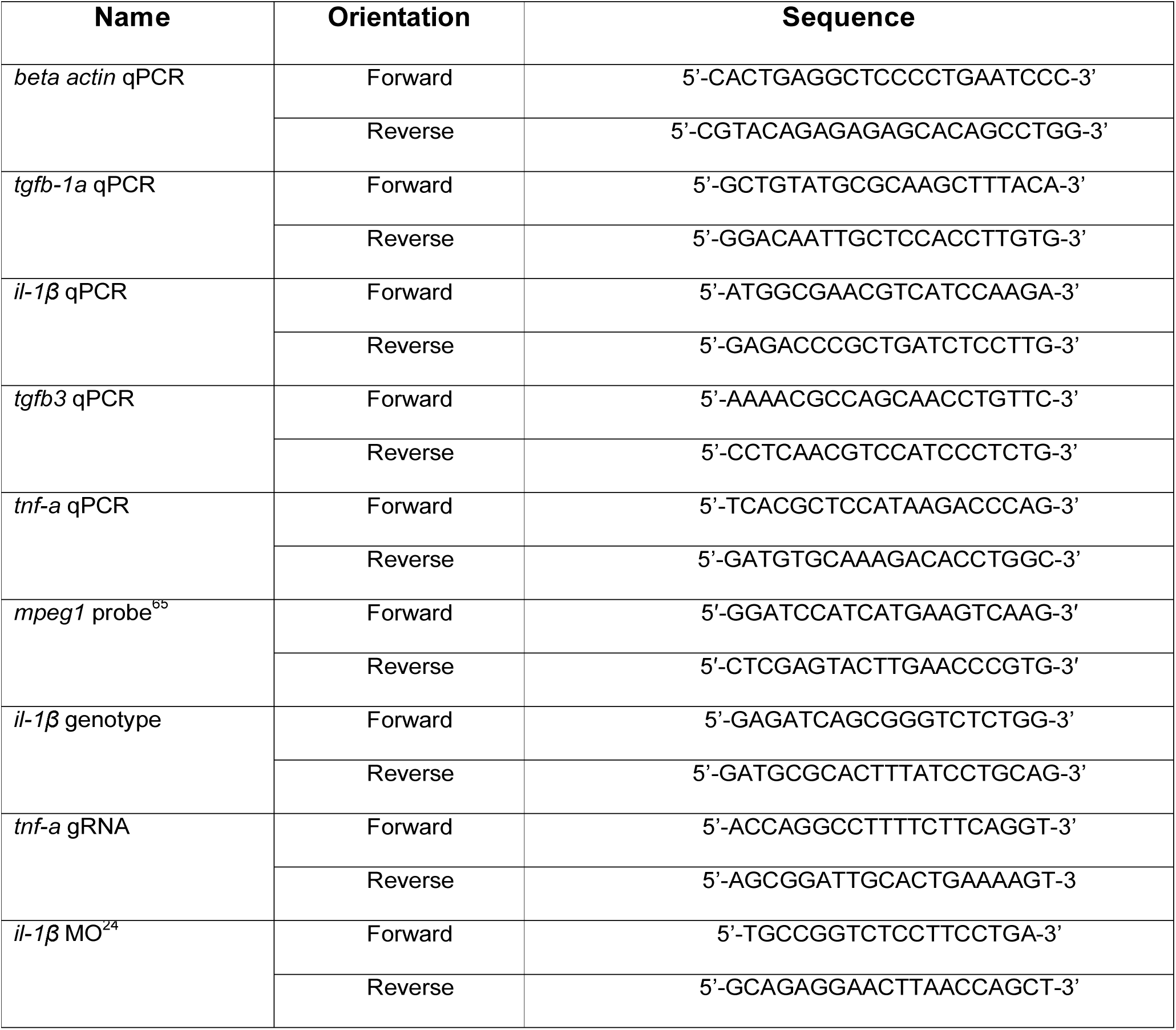
List of primers

### In situ Hybridization

Whole mount in situ hybridization was performed as previously described ^1^. Briefly, after fixation in 4% PFA, larvae were digested with 40μg/ml Proteinase K (Invitrogen). Thereafter, larvae were washed briefly in PBT and were re-fixed for 20 min in 4% PFA followed by washes in PBT. After washes, larvae were incubated at 67°C for 2h in pre-warmed hybridization buffer. Hybridization buffer was replaced with digoxigenin (DIG) labelled ISH probes diluted in hybridization buffer and incubated at 67°C overnight. The next day, larvae were washed thoroughly at 67°C with hybridization buffer, 50% 2×SSCT/50% deionized formamide, 2xSSCT and 0.2xSSCT. Larvae were then washed in PBT and incubated for 1 h in blocking buffer under slow agitation. Thereafter, larvae were incubated overnight at 4°C in blocking buffer containing pre-absorbed anti-DIG antibody. The next day, larvae were washed in PBT, followed by washes in staining buffer. Colour reaction was performed by incubating larvae in staining buffer supplemented with NBT/BCIP (Sigma-Aldrich) substrate. The staining reaction was terminated by washing larvae in PBT.

### Immunofluorescence

All incubations were performed at room temperature unless stated otherwise. For most immunolabelling experiments, the larvae were fixed in 4% PFA-PBS containing 1% DMSO at 4°C overnight. After washes in PBS, larvae were washed in PBTx. After permeabilization by incubation in PBS containing 2 mg/ml Collagenase (Sigma) for 25 min larvae were washed in PBTx. They were then incubated in blocking buffer for 2 h and incubated with primary antibody (1:50–1:500) diluted in blocking buffer at 4°C overnight. On the following day, larvae were washed times in PBTx, followed by incubation with secondary antibody diluted in blocking buffer (1:300) at 4°C overnight. The next day, larvae were washed three times in PBTx and once in PBS for 15 min each, before mounting in glycerol.

For whole mount immunostaining using primary antibodies from the same host species (rabbit anti-Il-1b, rabbit anti-Mpx, rabbit anti-Tp63) the samples were treated as previously described by others ^52^. Briefly, after fixation the samples were initially incubated with the first primary antibody at 4^°^C overnight. After washes with PBTx the samples were incubated with the conjugated first secondary antibody overnight at 4 ^°^C. Subsequently samples were incubated with blocking buffer for 1 h at RT in order to saturate open binding sites of the first primary antibody. Next, the samples were incubated with unconjugated Fab antibody against the host species of the primary antibody in order to cover the IgG sites of the first primary antibody, so that the second secondary antibody will not bind to it. After this, samples were incubated with the second primary antibody overnight at 4^°^C and subsequently with the second conjugated secondary antibody overnight at 4^°^C before mounting in glycerol. No signal was detected when the second primary antibody was omitted, indicating specificity of the consecutive immunolabeling protocol.

For whole mount immunostaining of acetylated tubulin, the protocol was performed as previously described ^1^. Briefly, larvae were fixed in 4% PFA for 1 h and then were dehydrated and transferred to 100% MeOH and then stored at - 20°C overnight. The next day head and tail were removed, and the samples were incubated in pre-chilled Acetone. Thereafter larvae were washed and digested with Proteinase K and re-fixed in 4% PFA. After washes the larvae were incubated with BSA in PBTx for 1 h. Subsequently the larvae were incubated for 2 overnights with primary antibody (acetylated tubulin). After washes and incubation with the secondary antibody the samples were washed in PBS for 15 min each, before mounting in glycerol.

### Evaluation of cell death using Acridine orange

In order to assess the levels of cell death after injury we used the acridine orange live staining as described by others ^53^. Briefly, at 1 and 2 dpl the larvae were incubated in 2.5 μg/ml solution of dye diluted into conditioned water for 20 minutes. After the staining, the larvae were washed by changing the water and larvae were live-mounted for imaging.

### Identification of dying cells after injury

In order to assess the levels of cell death after injury cross sections of larvae were used. Larvae were fixed in 4% PFA overnight at 4^°^C. After washes with 0.5% PBSTx, the larvae were transferred to 100% methanol and incubated for 10 min at room temperature. After rehydration, the larvae were washed with PBSTx 0,5%. Following this, larvae were mounted in 4% agarose and 50μm sections were performed using a vibratome (MICROM HM 650 V). The sections were then permeabilized using 14μg/ml diluted in 0,5%PBSTx. After brief wash with PBSTx the sections were postfixed in 4%PFA for 20min. Excess PFA was washed out and the samples were incubated with the TUNEL reaction mix according to the In-Situ Cell Death Detection Kit TMR red protocol (Roche).

### Morpholino injection

All morpholinos were injected into single cell stage larvae in total volume of 2 nl. Knockdown of *il-1β* was carried out using the antisense morpholino against *il-1β* (5D-CCCACAAACTGCAAAATATCAGCTT-3D) targeting the splice site between intron 2 and exon 3 according to ^24^. In order to block neutrophil development, we used the previously described MO combination^15^ of *pu.1* (5’-GATATACTGATACTCCATTGGTGGT-3’)^54^ which targets the translational start (ATG) of the pu.1 and the splice blocking MO against *gcsfr* (5’-TTTGTCTTTACAGATCCGCCAGTTC-3’) ^55^. All morpholinos and standard control (5’-CCTCTTACCTCAGTTACAATTTATA-3’) were obtained from Gene Tools, LLC.

### CRISPR-mediated genome editing

CRISPR gRNA for *il-1β* was designed using CRISPR Design (http://crispr.mit.edu) and ZiFit (http://zifit.partners.org/ZiFiT) webtools. Vectors were made by ligating the annealed oligonucleotides into pT7-gRNA expression vector as previously described ^1^. The gRNA was transcribed using the mMESSAGE mMACHINE T7 kit (Ambion) and assessed for size and quality on an electrophoresis gel. The injection mix consisted of 75 pg *il-1β* gRNA (target sequence: 5’-TGTGGAGCGGAGCCTTCGGCGGG-3’) and 150pg Cas9 RNA and was injected into single cell stage larvae. The CrRNA for the *Tnf-α* (target sequence: 5’-CCCGATGATGGCATTTATTTTGT-3’) and the tracrRNA were ordered from (Merck KGaA, Darmstadt). The injection mix included 1μl Tracer 250 ng /μl, 1 μl gRNA, 1μl, 1μl Cas9 protein, 1μl RNAse free H20, 1μl fluorescent dextran. Larvae injected with GFP gRNA (target sequence: 5’GGCGAGGGCGATGCCACCTA-3) and uninjected larvae were used as controls. The efficiency of the mutagenesis was assessed by RFLP analysis.

### Live imaging of zebrafish larvae and time-lapse imaging

For the acquisition of all fluorescent images, LSM 710 and LSM 880 confocal microscopes were used. For live confocal imaging, zebrafish larvae were anesthetized in PBS containing 0.02% MS222 and mounted in 1.5% low melting point agarose (Ultra-PureTM, Invitrogen). During imaging, the larvae were covered with 0.01% MS222-containing fish water to keep preparations from drying out. For time-lapse imaging, agarose covering the lesion site was gently removed after gelation. Time-lapse imaging was performed for 19 h starting at 6 hpl. Acquired time-lapse images were denoised using the ImageJ plugin CANDLE-J algorithm, which allows removal of the noise compartment while preserving the structural information with high fidelity ^56^. Comparison of raw movies with CANDLE-J-processed movies showed that edges of features remained conserved after denoising.

### Scoring of spinal cord bridging

Axonal bridging was scored in fixed and live mounted samples using fluorescence-equipped stereomicroscope (Leica MDG41) and confocal microscopes (LSM 710, LSM 880) at time points of interest as previously described ^1^. For live analysis of transgenic larvae, the observer was blinded to the experimental condition before scoring. For confocal analysis, scoring of images was performed blinded to the experimental condition. All analyses were performed blinded to the experimental condition on at least three independent clutches of larvae.

### Cell counting in whole-mounted larvae

A volume of interest was defined centered on the lesion site from confocal images. The dimensions were: width = 200 μm, height = 75 μm (above the notochord), depth = 50 μm. Images were analysed using the Imaris (Bitplane, Belfast, UK) or the ImageJ software. The number of cells was quantified manually in 3D view, blinded to the experimental condition on at least three independent clutches of larvae.

### Quantification of diffuse signals in whole-mount larvae

For most quantifications of diffuse signal (Fig.4B and Fig.S1), image stacks, centered in the lesion site (height: 50 μm) were collapsed and an area of interest defined: width = 100 μm, height = 50 μm (above the notochord). The image was thresholded and the ‘’Analyze Particles’’ tool in Fiji ^57^ with default settings was used to calculate the percentage of area taken up by signal.

To quantify Col I in the injury site (Fig. S3C) we used a previously published protocol ^1^. Briefly, image stacks were collapsed, thresholded and the area of the signal was measured.

All analyses were performed blinded to the experimental condition on at least three independent clutches of larvae.

### Statistical analysis

Power analysis using G*Power ^58^, was used to calculate power (aim > 0.8) for the experiments and determine the group sizes accordingly. Statistical power was > 0.8 for all experiments. All quantitative data were tested for normality and analyzed with parametric and non-parametric tests as appropriate. The statistical analysis was performed using IBM SPSS Statistics Shapiro-Wilk’s W-test was used in order to assess the normality of the data. Quantitative RT-PCR data were analyzed as previously described ^59,60^. Kruskal-Wallis test followed by Dunn’s multiple comparisons, One-way ANOVA followed by Bonferroni multiple comparisons test, two-way ANOVA, followed by Bonferroni multiple comparisons, t-test, Mann–Whitney U test or Fischer’s exact test were used, as indicated in the Figure legends. *P< 0.05, **P< 0.01, ***P< 0.001, n.s. indicates no significant. Error bars indicate the standard error of the mean (SEM). The Figures were prepared with Adobe Photoshop CC and Adobe Illustrator CC. Graphs were generated using GraphPad Prism 7.

## ACKNOWLEDGEMENTS

We thank Dr. William Talbot and Dr. Stephen Renshaw for sharing mutants before publication, Dr. Bertrand Vernay for expert advice on microscopy and Joli Ghanawi for excellent fish facility management. Supported by the BBSRC to CGB and TB, NC3Rs PhD studentship to CGB/TB for TMT, DFG to DW, Summer research studentships to TM (Carnegie Trust) and EK (Anatomical Society). NO was funded by Biotechnology and Biological Sciences Research Council (BBSRC) project grant (BB/L000830/1).

## AUTHOR CONTRIBUTIONS (CRediT standard)

Conceptualisation – TMT, TBe, CGB; Investigation – TMT, DW, LC, TM, MK, ML, AU, TBa, EK; Resources – NO, SAR, YF, TJH; Supervision – DW, NO, YF, TBe, CGB; Writing – TMT, TBe, CGB; Funding Acquisition – TBe, CGB

---

**Fig. S1:**
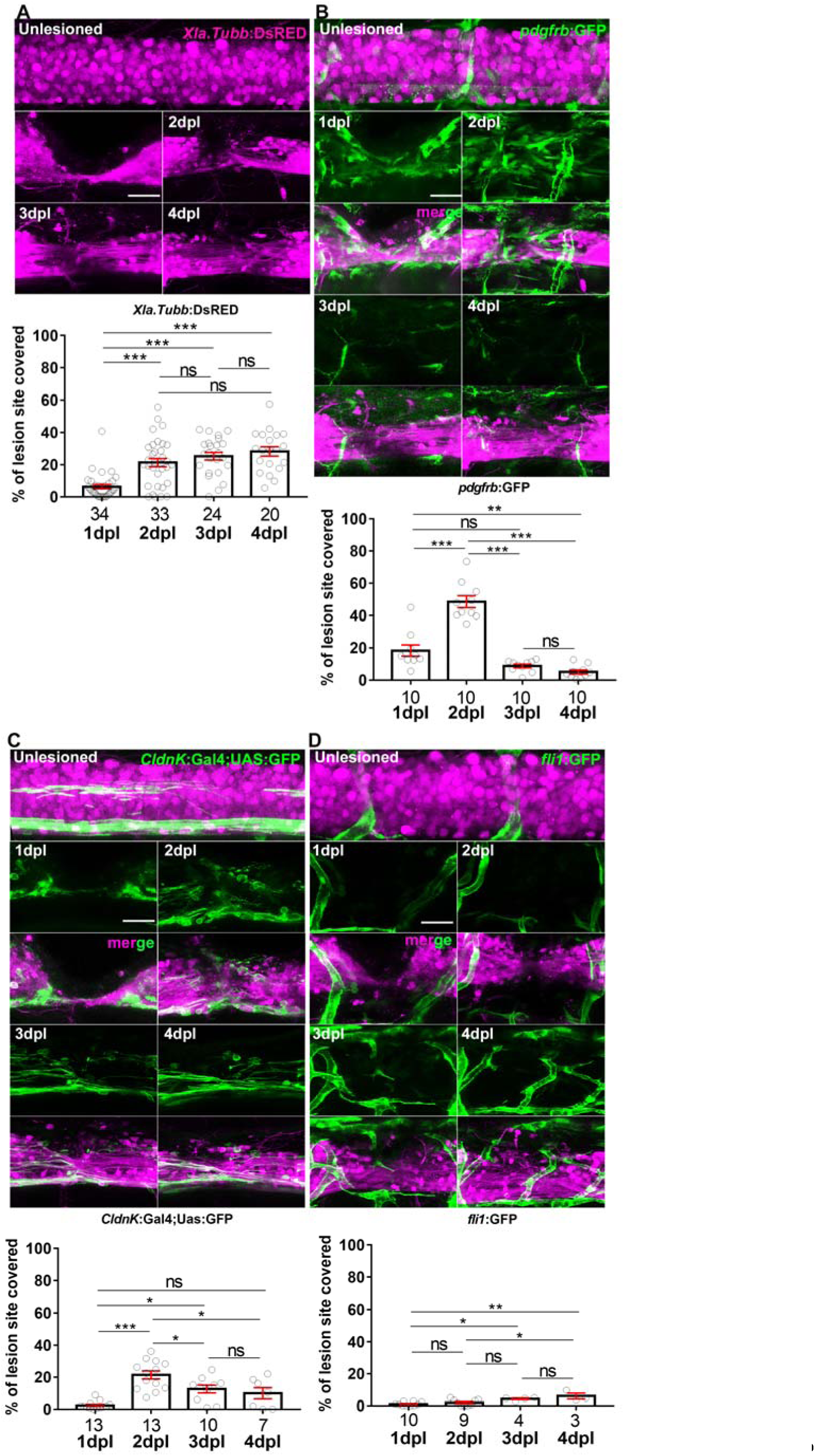
Presence of fibroblast-like cells, but not myelinated cells or endothelial cells correlates with axonal regeneration. **A:** Axonal density in *XIa.tubb*:DsRed transgenic animals increases after injury starting at 1 dpl and plateauing at 2 dpl (*One-way ANOVA followed by Bonferroni multiple comparisons:* F _3, 107_ = 19.67, ***P < 0.0001, ns indicates no significance). **B:** Presence of fibroblast-like cells (*pdfrb*:GFP^+^) in the lesion site is substantial from 1 dpl, it peaks at 2 dpl, and declines thereafter (*One-way ANOVA followed by Bonferroni multiple comparisons:* F _3, 36_ = 53.92, **P<0.01, ***P<0.0001, n.s indicates no significance). **C:** Myelinating cells (*cldnK*:GFP^+^) appear in the injury site only at 2 dpl (*One-way ANOVA followed by Bonferroni multiple comparisons:* F _3, 39_ = 14.52, *P0.05, ***P<0.0001, n.s indicates no significance). **D:** Endothelial cells (*fli1*:GFP^+^) do not invade the lesion site (*One-way ANOVA followed by Bonferroni multiple comparisons*: F _3, 22_ = 7.939, *P0.05, **P<0.01). Identical animals are imaged on consecutive days. Lateral views of the injury site are shown; rostral is left. *Scale bar:* 25 μm. *Error bars* indicate SEM.

**Fig. S2:**
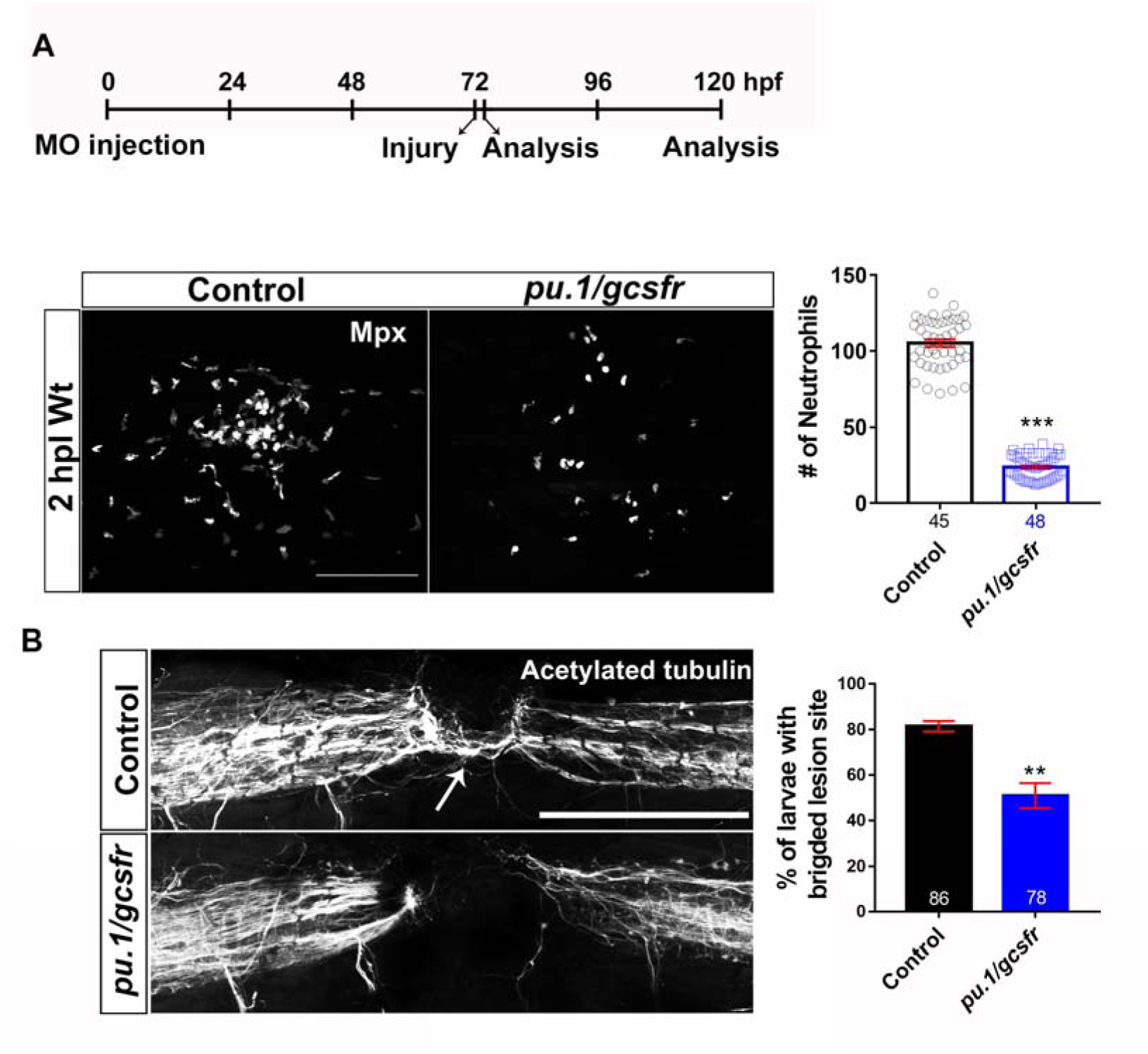
Knock down of immune cell development impairs axonal regeneration. **A:** Neutrophils are strongly reduced in number in *pu.1* and *gcsfr* double morpholino-injected animals (*Mann-Whitney test*: ***P<0.001). **B:** Quantification of axonal bridging (arrow; anti-acetylated Tubulin) shows that morpholino injection decreases the proportion of animals with axonal bridges (*Fisher’s exact test*: **P<0.01). Lateral views of the injury site are shown; rostral is left. *Scale bar:* 100 μm. *Error bars* indicate SEM.

**Fig. S3:**
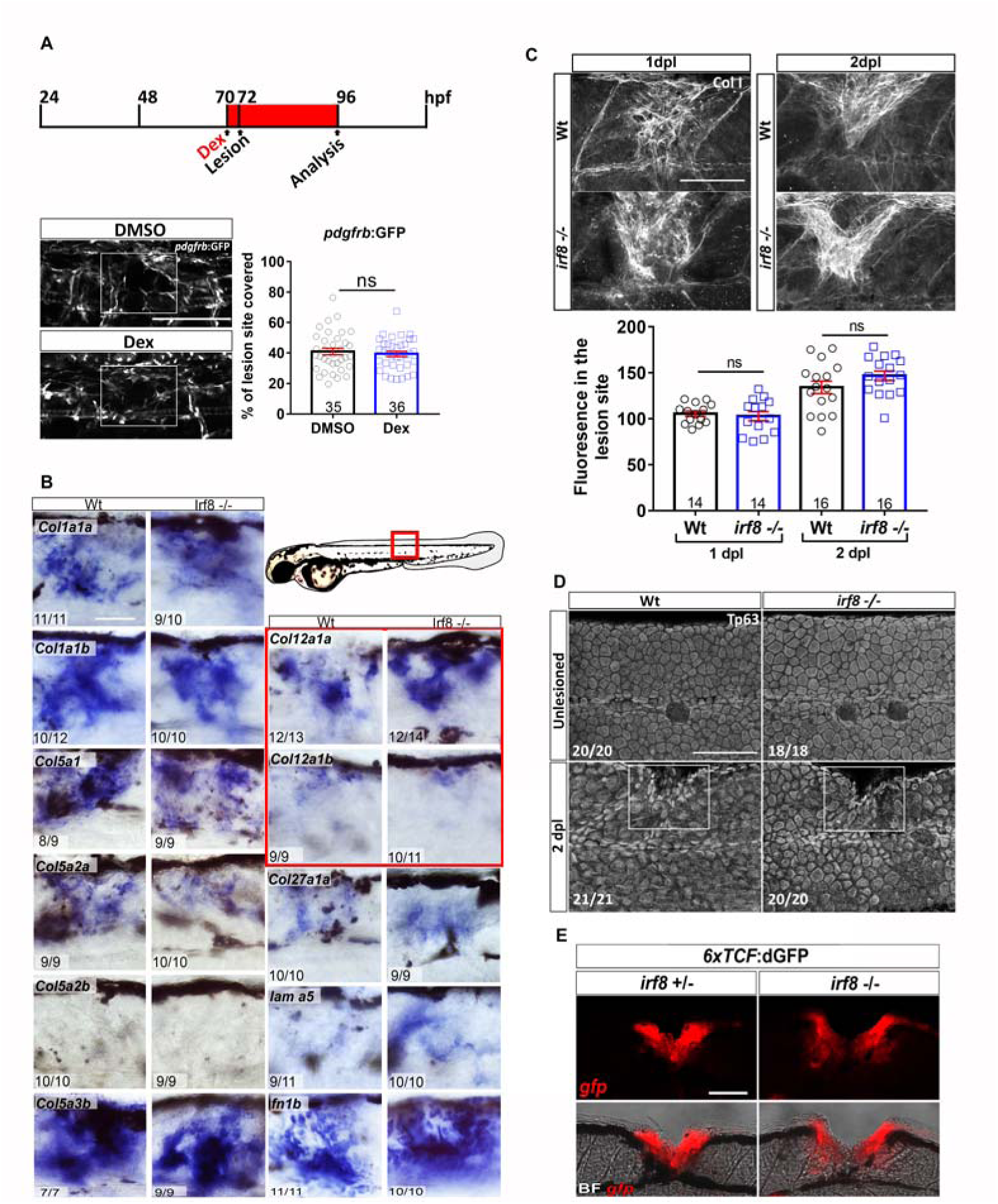
Wnt-dependent ECM deposition and wound healing are not affected by compromising the immune reaction. Lateral views of the injury site are shown.; rostral is left. **A:** The density of fibroblast-like cells (*pdgfrb*:GFP^+^) in the lesion site is not affected by inhibition of the immune response with dexamethasone (Dex) (*t-test*, n.s. = no significance). **B:** Expression of mRNAs for major ECM components is not impaired in the absence of macrophages in *irf8* mutants. **C:** Immunoreactivity for Collagen I (Col I) is not altered in *irf8* mutants, compared to wildtype controls at 1dpl and 2dpl (*t-test*: n.s indicated no significance). **D:** Basal keratinocytes (anti-Tp63^+^) cover the injury site (indicated by rectangle) in *irf8* mutants by 2dpl, similar to wildtype animals. **E:** Activation of the Wnt pathway is unaltered in *irf8* mutants after injury. The *6xTCF*:dGFP reporter line crossed into the *irf8* mutants shows activation of Wnt signalling that is comparable to that in wildtype animals at 2 dpl. Error bars indicate SEM. Scale bars: 50μm.

**Fig. S4:**
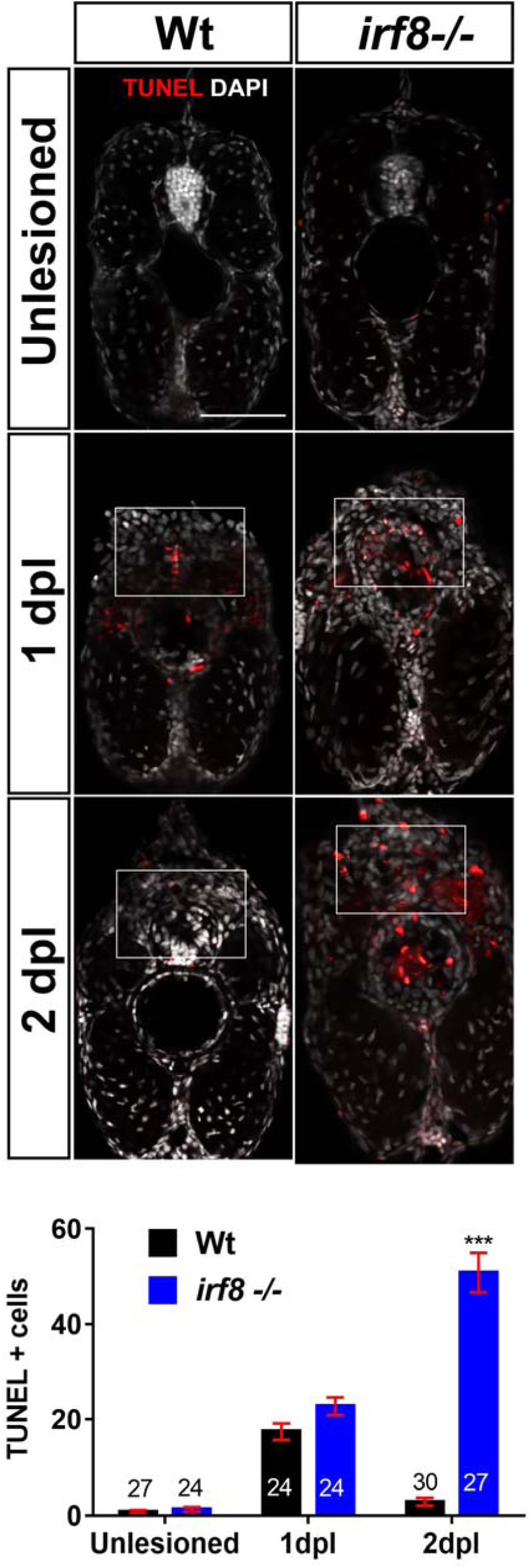
The number of dying cells is strongly increased in the lesion site of *irf8* mutants at late stages of regeneration. TUNEL labelling on cross section of larvae (dorsal is up) shows an increased number of TUNEL^+^ cells in the injury site (indicated by rectangle) of the *irf8* mutant fish at 2 dpl, but not at 1 dpl (*t-test*: ***P<0.001, n.s = no significance). *Scale bar:* 25 μm. *Error bars* indicate SEM.

**Fig. S5:**
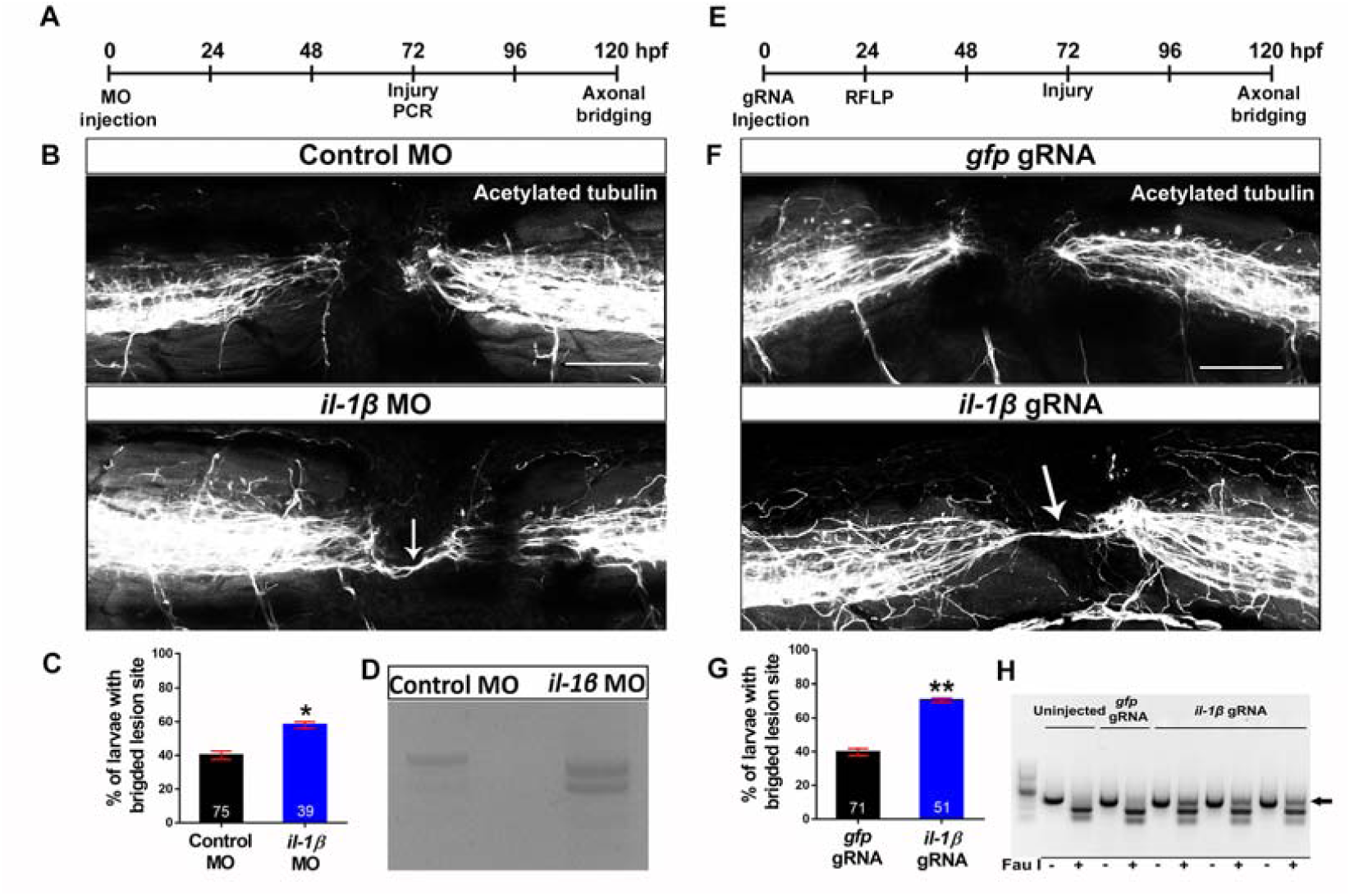
Genetic inhibition of *il-1β* function rescues axon bridging in *irf8* mutants. Lateral views of the injury site are shown; rostral is left. **A:** The timeline for the morpholino manipulations is given. **B-D:** Morpholino gene knockdown of *il-1β* by injecting into the zygote shows characteristic mis-splicing of *il-1β* mRNA, compared to wildtype animals (D) and rescues the bridging phenotype of the *irf8* mutant fish (B,C; *Fisher’s exact test*: *P<0.05). **D:** Morpholino gene knockdown of *il-1β* shows characteristic mis-splicing of *il-1β* mRNA, compared to wildtype animals. **E:** The timeline for gRNA manipulation is given. **F-G:** CRISPR/Cas9-mediated disruption of *il-1β* by injecting specific gRNA into the zygote, leads to somatic mutations, as shown by RFLP analysis (arrow in **H**) and rescues the bridging phenotype in *irf8* mutants (F,G; *Fisher’s exact test*: **P<0.01). Error bars represent SEM. Scale bars: 50 μm.

**Fig. S6:**
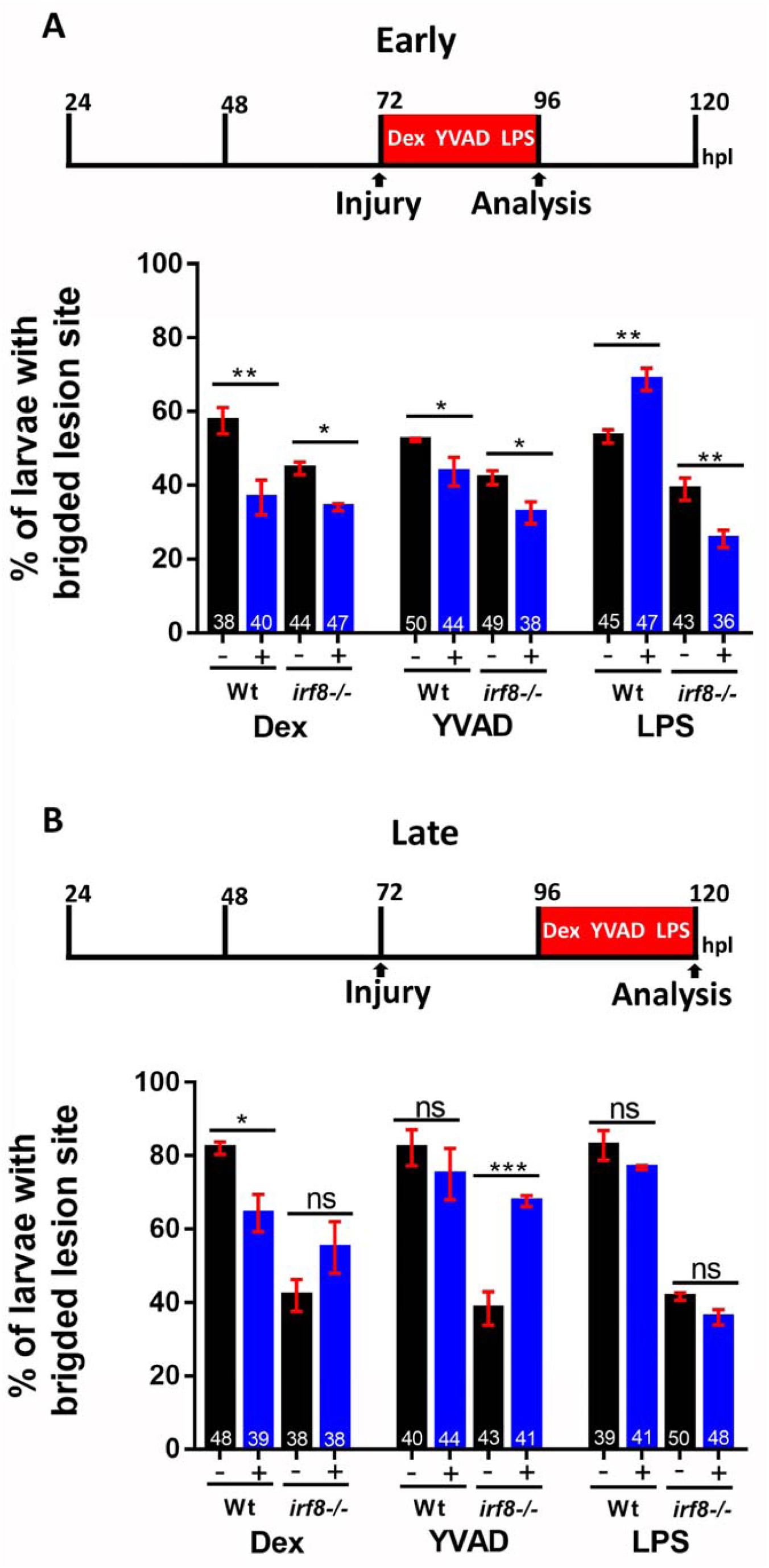
Temporally restricted manipulations of the immune response reveal time-dependent effects on axon bridging. **A:** During early regeneration, dexamethasone and YVAD treatment impair regeneration in both wildtype animals and *irf8* mutants. Stimulation of the immune response with LPS promotes regeneration in wildtype animals but inhibits it in *irf8* mutants (*Two-way ANOVA followed by Bonferroni multiple comparisons*: F _5, 12_ = 25.32. *P<0.05, **P<0.01). **B:** Manipulations during late regeneration show persistent negative effects of dexamethasone in wildtype animals and a strong rescuing effect of YVAD in *irf8* mutants (Two-way ANOVA followed by Bonferroni multiple comparisons: F _5, 12_ = 16.05. *P<0.05, ***P<0.001, n.s. = no significance. Error bars represent SEM.

**Fig. S7:**
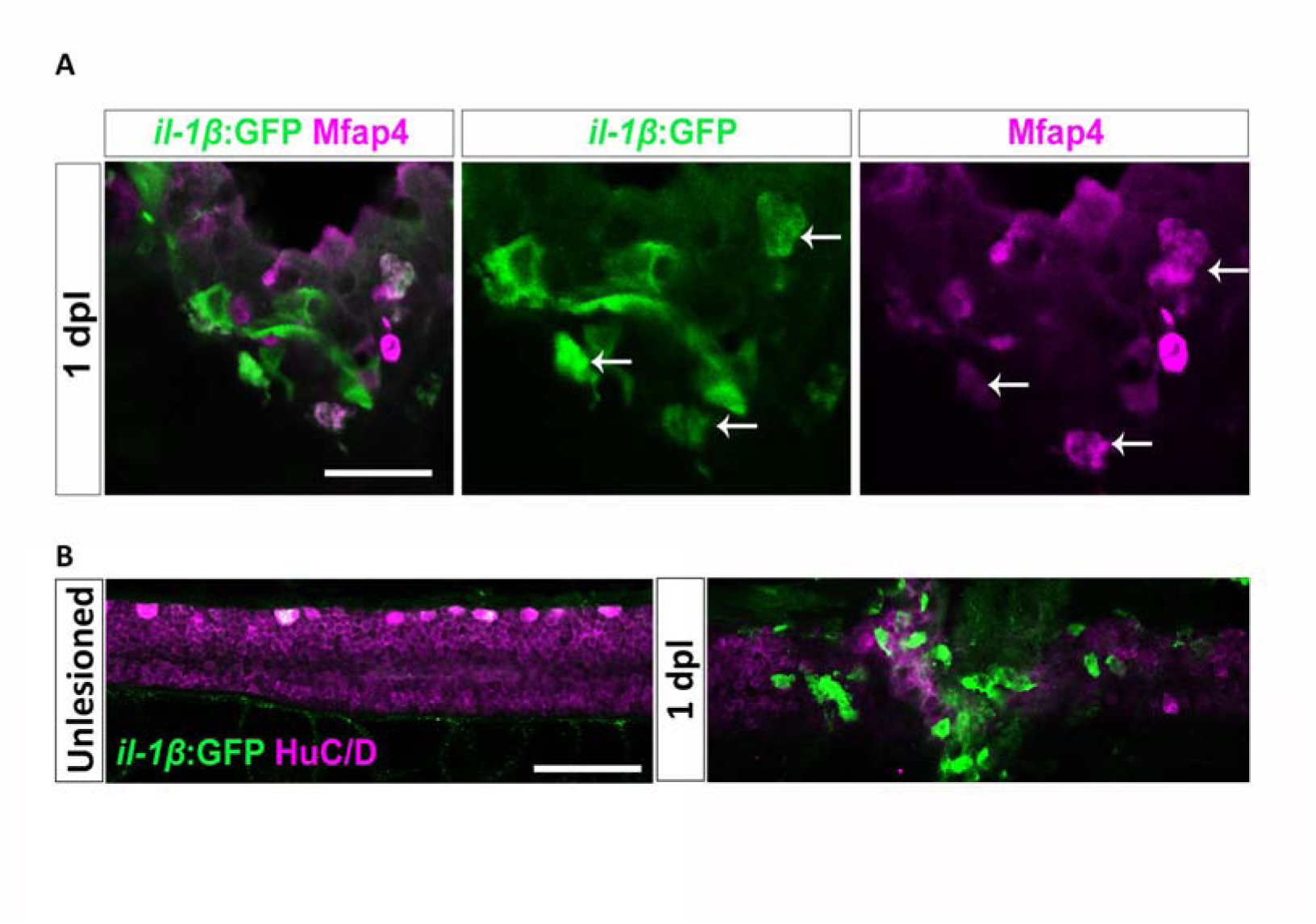
*il-1β* is expressed in macrophages, but not neurons in wildtype animals. **A:** In the il-1β reporter line, *il-1β*:GFP^+^ cells were frequently co-labelled (arrows) with the macrophage marker Mfap4 at 1 dpl. **B:** *il-1β*:GFP^+^ cells were rarely co-labelled with the neuronal marker HuC/D at 1 dpl. Single optical sections are shown. Scale bars: 50 μm.

## List of Movies

Movie S1: mpeg1:mcherry;mpx:gfp double transgenic animals, showing the recruitment of macrophages and neutrophils to the injury site.

Movie S2: Xla.Tubb:DsRed;mpeg1:EGFP double transgenic animals, showing macrophage phagocytosis of neuronal debris and axonal bridging with no interaction between immune cells and axons.

